# Fungal community composition links rhizosphere microbiome organization to plant phenotype in response to moderate warming

**DOI:** 10.64898/2026.05.29.728675

**Authors:** Xin Li, Jana Trenner, Lennart Eschen-Lippold, John S. Park, Alexandra Boritzki, Mika Tarkka, Kirsten Küsel, Christine Römermann, Ivo Grosse, Stéphane Hacquard, Jon Ågren, Carlos Alonso-Blanco, Jun Zhou, Marcel Quint

## Abstract

**Background:** Global warming increasingly challenges plant performance and ecosystem function, and plant responses to elevated temperature are shaped not only by intrinsic plasticity, but also by interactions with rhizosphere microbial communities. However, it remains unclear how warming-induced microbiome reorganization relates to plant performance, whether microbiome community composition predicts plant phenotypic responses to elevated temperature better than simple host-microbiome origin matching, and which bacterial or fungal community features contribute most strongly to this prediction.

**Results:** We implemented a full factorial design manipulating plant genotype (P), microbial inoculum (M), and temperature (T) using natural *Arabidopsis thaliana* ecotypes and their corresponding rhizosphere microbiomes from contrasting climatic regions. By integrating high-throughput plant phenotyping, microbial community profiling, microbiome-phenotype coupling analyses, and predictive modeling, we found that elevated temperature induced a coherent thermomorphogenic shift in plant architecture, while plant phenotypic variation remained determined primarily by genotype over the 14-day vegetative growth period covered in this study. Overall, cold-origin genotypes showed larger temperature-induced trait shifts than warm-origin genotypes. Rhizosphere microbial communities were structured predominantly by inoculum origin, but were also affected by warming. Evidence for a increased thermomorphogenic growth when genotypes were combined with inoculum from their home site was limited. Instead, microbiome-phenotype associations were better captured by community composition than by matching status, with fungal community variation showing stronger and more consistent associations with plant phenotype than bacterial variation and providing more robust predictions of plant phenotypes across models.

**Conclusions:** Warming reorganized plant-rhizosphere systems within persistent host- and inoculum-associated baselines. Plant phenotypic variation during vegetative development under elevated temperature was linked more closely to microbiome composition, especially in fungi, than to simple host-microbiome matching of origins. These findings provide a framework for identifying microbiome features associated with plant performance under climate warming.

## BACKGROUND

Rising global temperatures are reshaping ecological systems from organismal physiology to community organization [1]. Climate projections indicate that plants will increasingly experience thermal regimes exceeding historical baselines [2], with consequences for species distributions, biodiversity patterns, and agricultural productivity. Plant responses to warming, however, are embedded in ecological networks rather than expressed in isolation. In terrestrial ecosystems, these networks include plant-associated microbial communities that interact closely with roots and influence plant nutrition, development, and stress tolerance [3–5]. Microbial communities are themselves shaped by plant hosts, environmental conditions, and interactions among microorganisms. Understanding how these biotic and abiotic forces jointly influence microbiome-mediated plant performance is therefore essential for predicting ecological responses to climate warming and for maintaining ecosystem functioning under rising temperatures [3].

Plants are sessile organisms that must continuously integrate environmental signals into developmental programs to sustain growth and reproduction. In the model plant *Arabidopsis thaliana*, moderately elevated ambient temperatures (approximately 26-28 °C) induce coordinated morphological changes, including petiole elongation, hyponastic growth (upward bending of leaves), and a more open rosette architecture [6,7]. These responses, collectively termed thermomorphogenesis, reflect a reallocation of growth that helps maintain physiological performance under warm conditions [6,8,9]. Although the molecular basis of thermomorphogenesis has been intensively studied, plant responses to warming are not determined by plant physiology alone. Root-associated microbial communities can influence plant growth through nutrient mobilization, hormone modulation, and protection against environmental stress [10–13]. Temperature-driven changes in rhizosphere microbial communities may therefore interact with plant thermomorphogenic responses and shape plant architecture under elevated temperature [3,14].

The rhizosphere microbiome is structured by multiple interacting drivers, including plant genotype, microbial source environment, and abiotic conditions such as temperature [15–18]. In many systems, soil origin explains a substantial proportion of microbial community variation, establishing a baseline upon which host filtering and environmental perturbations act [17,19]. Large-scale surveys of *A. thaliana* across Europe have provided important insight into these assembly processes. Continental-scale analyses revealed strong geographic structuring of soil microbial communities, reflecting environmental heterogeneity across sites, whereas root-associated bacterial assemblages showed partial convergence across locations, suggesting that a limited set of phylogenetically diverse bacteria repeatedly colonizes plant roots despite differences in surrounding soils [17]. Reciprocal transplant experiments between geographically distant *A. thaliana* populations further showed that plant adaptation across climatic gradients is driven primarily by local environmental conditions, while soil microbial communities contribute comparatively little to the magnitude of adaptive differentiation [16]. These studies highlight the joint roles of host genotype, environmental context, and microbiome assembly in shaping plant-microbiome systems.

Temperature is a fundamental regulator of microbial metabolism and growth [16–18]. Warming experiments across ecosystems have documented shifts in microbial community composition and associated ecosystem processes under elevated temperature [20–22]. Microbial taxa often occupy distinct climatic niches, and warming can therefore drive taxon-specific changes in microbial richness and phylogenetic structure [23]. Plant-associated fungal communities frequently show stronger compositional responses to elevated temperature than bacterial communities, pointing to potential kingdom-level differences in sensitivity to thermal perturbation [24,25]. Beyond taxonomic composition, warming may also alter the ecological organization of microbial communities by changing interaction patterns and overall community structure. Such changes can influence the balance between cooperative and competitive associations and may affect microbiome stability. Because plant-associated microbiomes regulate processes central to plant performance, including nutrient acquisition, hormone signaling, and stress tolerance, warming-induced reorganization of plant-microbe interactions has the potential to affect both plant phenotype and ecosystem functioning.

Despite growing recognition that plant-associated microbiomes influence plant performance, the mechanisms linking warming-induced microbiome reorganization to plant phenotypic responses remain poorly resolved. It remains unclear whether elevated temperature overrides host- and soil-origin effects on rhizosphere community composition or instead adds temperature-dependent shifts while preserving differences associated with plant genotype and soil inoculum origin. It is also unclear whether plant responses to warming are equally strongly associated with bacterial and fungal community composition.

To address these questions, we implemented a fully factorial design manipulating plant genotype (P), microbial inoculum (M), and temperature (T) using natural *Arabidopsis thaliana* ecotypes and their corresponding rhizosphere microbiomes from contrasting climatic regions. We used 16 °C as a cool reference condition and 28 °C as a warm treatment, thereby establishing a defined thermal contrast between a lower baseline temperature and a widely used thermomorphogenic setpoint. This design allowed us to examine microbiome-associated plant responses under controlled warming conditions while maintaining comparability with the *Arabidopsis* thermomorphogenesis literature. Specifically, we aimed to (i) determine how elevated temperature reshapes plant morphology and rhizosphere microbial community structure; (ii) test whether changes in microbiome composition, rather than simple host-microbiome matching by origin, helps explain variation in plant performance under warming; and (iii) identify which aspects of microbiome composition are the best predictors of plant phenotypic variation under elevated temperature, and whether correlations with bacterial and fungal community compositions differ in strength. By integrating plant phenotyping, microbial community profiling, correlation analyses between microbiome composition and plant phenotypes, and predictive modeling, this study links thermal perturbation to the compositional organization of rhizosphere microbiomes and provides a framework for understanding how plant-microbiome systems reorganize under climate warming.

## METHODS

### *A. thaliana* source populations and soil collection

To guide the selection of source populations for the factorial experiment, we first carried out a two-step pilot screen using European *A. thaliana* accessions from the 1001 Genomes seed collection [26]. In the first step, 114 accessions from habitats spanning the warm and cold margins of the European climatic range of *A. thaliana* were grown on agar plates for 7 days, and hypocotyl responses under warm versus cool ambient conditions were quantified. Based on this screen, 48 accessions showing pronounced differences in temperature-responsive growth, together with *Col-0* (1001g ID: 6909) as a reference genotype, were selected for a second pilot experiment in soil. This follow-up assay was performed under the same cultivation conditions as the main experiment, and plants were imaged on day 11 of the temperature treatment to quantify early vegetative traits. The soil-based screen was used to prioritize sampling regions that combined clear temperature-response contrasts with warm versus cold climatic origins and broadly comparable site conditions, particularly soil pH, carbon/nitrogen status, and habitat context. Six field sites were therefore selected: three in central Spain and three in Sweden, representing warm and cold extremes of the European *A. thaliana* range while minimizing major site-level differences where possible (**Table S1**).

Up to 8 plants were collected in March (Spain) and May (Sweden) for each site at the same developmental stage (bolting/flowering stage; **Table S1**). To minimize disturbance to the root system, plants were harvested together with the surrounding bulk soil and then transferred to pots in the greenhouse. Plants were maintained under controlled greenhouse conditions for seed production to rule out residual maternal environmental effects that might otherwise interfere with genotype effects in downstream analyses. Greenhouse-propagated seeds of several mother plants were bulked and used in the reciprocal planting experiment.

For soil collection at the exact plant sampling sites, the uppermost ca. 5 cm were discarded, and soil samples were taken from a depth of ca. 5-25 cm. Soils were air-dried at room temperature for less than 2 weeks and then stored in the dark at 4 °C until use.

### Soil microbiome transplants

To minimize differences in soil matrix while preserving source-specific microbial inocula, sterilized recipient soil (peat soil from PATZER Erden ORANGE VM Tr) was thoroughly mixed with donor soil collected from each site, which served as the microbial inoculum (M), at a ratio of 9:1 (recipient soil: donor soil). Sterility of the peat soil was verified by plating soil suspensions on 50% Tryptic Soy Agar (TSA) and incubating them at 25 °C for up to 7 days. After inoculation, the soils were incubated at room temperature for 2 weeks to allow microbial acclimation before seedling transplantation [16]. Repopulated soils were then collected for soil property measurements and amplicon sequencing.

### Experimental treatments and growth conditions

To investigate the three-way interaction among plant genotype (P), microbial inoculum (M), and temperature (T), a full factorial design (P × M × T) was established. Seven *A. thaliana* genotypes were used: three from Spain, three from Sweden, and the reference genotype *Col-0* (1001g ID: 6909). These were grown in seven soil treatments: three carrying Spanish microbial inocula, three carrying Swedish microbial inocula, and one sterilized-soil control, under 16 °C and 28 °C conditions. The experimental design is shown in **Fig. 1**. Climatic background data for the collection sites were obtained from CHELSA-derived environmental layers compiled in AraCLIM V2.0 within Arabidopsis CLIMtools [27]. We used the CHELSA spring mean temperature (CHELSA Tmean spring) and spring maximum temperature (CHELSA Tmax spring) variables to describe long-term temperature conditions at accession origins, because spring better approximates the main vegetative growth window of *A. thaliana* in its natural habitats. Although mean seasonal maximum temperatures at the collection sites remained below the 28°C setpoint (**Table S1**), the 16-28 °C temperature regime was chosen to a) maintain comparability with the *Arabidopsis* thermomorphogenesis literature, and to b) ensure a significant extent of thermomorphogenic plant growth behaviour. As such, 28 °C represent an upper-end warming scenario rather than a simulation of average spring temperatures in the field.

**Fig. 1.**
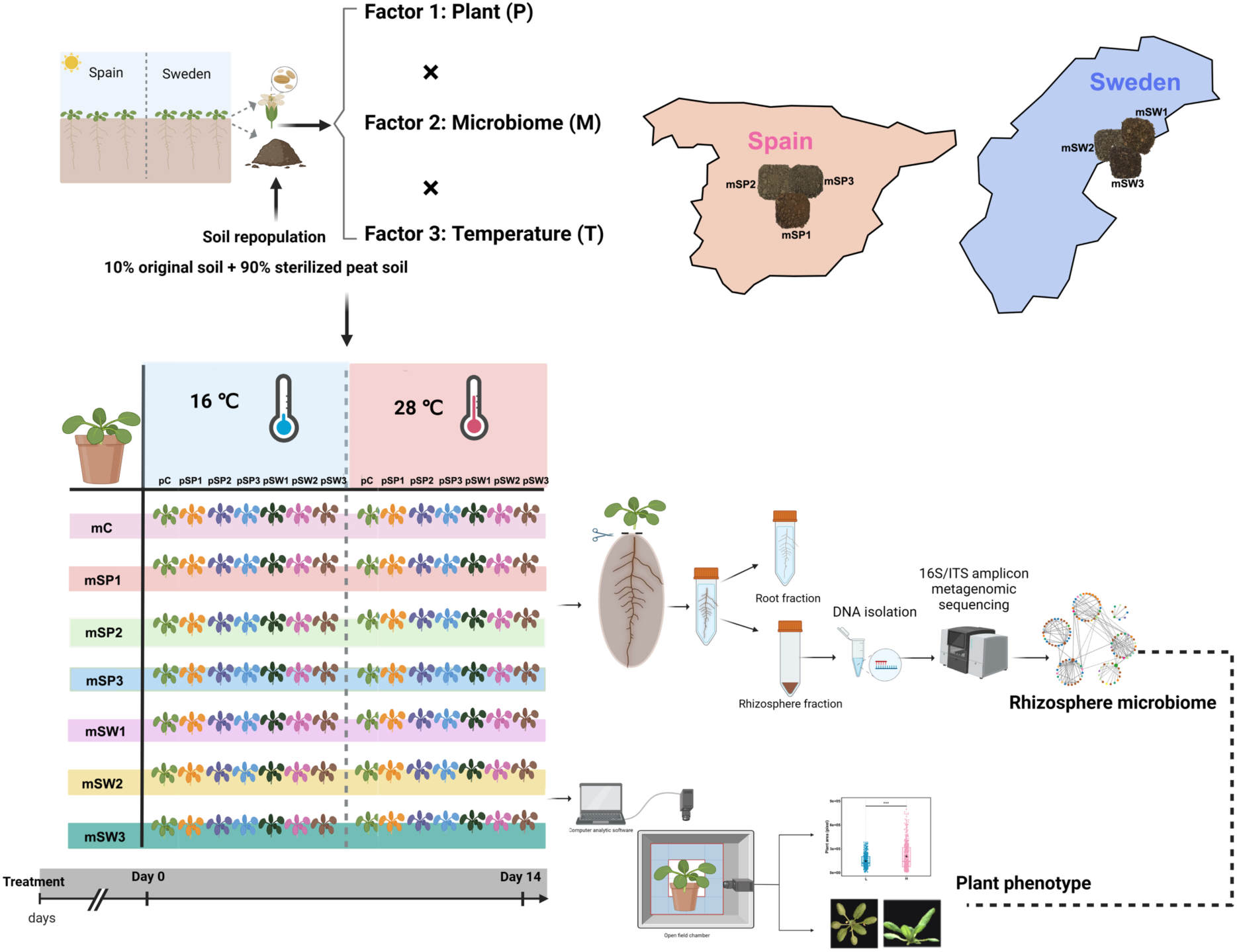
Experimental design of the *Arabidopsis*-rhizosphere microbiome reciprocal inoculation experiment. Natural *A. thaliana* genotypes from warm (Spain) and cold (Sweden) climatic regions were grown with microbial inocula derived from rhizosphere soils collected at the corresponding field sites. A full factorial design manipulated plant genotype (P), microbial inoculum (M), and temperature (T), with plants cultivated under 16 °C and 28 °C conditions. Temperature levels were 16 °C (L) and 28 °C (H). Microbial inoculum treatments included sterilized peat soil control (mC), Spanish inocula (mSP1-mSP3), and Swedish inocula (mSW1-mSW3). Plant genotypes included the control genotype (pC), Spanish genotypes (pSP1-pSP3), and Swedish genotypes (pSW1-pSW3). High-throughput phenotyping was conducted using the LemnaTec PhenoAIxpert platform (LemnaTec GmbH, Aachen, Germany). Image acquisition and trait extraction were performed using LemnaTec software (LemnaGrid, LemnaExperiment, Analysis Executor4, and LemnaAIxplorer). Rhizosphere bacterial and fungal communities were profiled by amplicon sequencing of the 16S rRNA gene and fungal ITS region. The reference genotype *Col-0* (1001g ID: 6909) and a sterilized-soil control were included in the experiment as additional controls.

Throughout the manuscript, plant genotypes and microbial inocula were grouped by climatic origin as warm (Spain) or cold (Sweden) where appropriate. For the primary home–away analysis, only the exact pairing between a plant accession and its native soil inoculum was classified as home. All other non-native combinations, including pairings within the same climatic region but from different sites, were classified as away. Col-0 and the sterilized-soil control were excluded from home-away analyses. The reference genotype *Col-0* and the sterilized-soil controls were included in the factorial experiment but were excluded from home-away matching analyses because they did not belong to either climatic origin group.

Seeds were surface-sterilized, rinsed thoroughly with sterile distilled water, and stratified in deionized water at 4 °C for 3 days before sowing. Seedlings were germinated on vertically oriented sterile agar plates containing solid *A. thaliana* solution (ATS) [28] supplemented with 1% (w/v) sucrose. Seedlings were grown for 14 days under long-day conditions (16 h light/8 h dark) at 16 °C with photosynthetically active radiation (PAR) of 90 μmol m⁻² s⁻¹ provided by white fluorescent lamps (T5, 4000 K) [29].

Uniformly germinated 14-day-old seedlings were transplanted into soil-filled pots and acclimated for 5 days in growth chambers set to 16/16 °C (day/night), 90 μmol m⁻² s⁻¹ light intensity, and a long-day photoperiod. Plants were then either maintained at 16/16 °C or shifted to 28/28 °C (day/night) for 14 days [30]. Temperature treatments were implemented in two chambers per setpoint, and environmental parameters (relative humidity, PAR, spectrum, temperature) were logged to ensure comparability across chambers (**Fig. S1**). Temperature treatments were implemented as fixed chamber setpoints. To reduce potential chamber- and position-related effects, pots were randomly repositioned and exchanged between the two chambers within each temperature treatment every second day. Each treatment combination consisted of 10 biological replicates. Differences between 16 °C and 28 °C are therefore interpreted as temperature-associated responses under controlled chamber conditions.

### Plant phenotypic trait recording and data analysis

High-throughput phenotyping was conducted using the LemnaTec PhenoAIxpert platform (LemnaTec GmbH, Aachen, Germany), a non-invasive RGB imaging system for automated monitoring of plant growth and morphology [31]. Image acquisition and trait extraction were performed using LemnaTec software (LemnaGrid, LemnaExperiment, Analysis Executor4, and LemnaAIxplorer). Plants were imaged from top and side views to quantify phenotypic traits. Day 0 refers to the onset of the temperature treatment, and the plant trait analyses reported here were based on endpoint measurements taken after 14 days of temperature treatment. Petiole length and petiole angle were additionally quantified from endpoint images using ImageJ. Fresh weight was also recorded at harvest. A total of 11 plant traits characterizing rosette size and shape were recorded (**Table S2**). In the main text, we focus on representative univariate endpoint traits and multivariate phenotype summaries. Here, plant performance refers to short-term vegetative performance assessed from endpoint traits after 14 days of temperature treatment, rather than lifetime fitness or growth rate.

To assess the effects of temperature, microbial inoculum, and plant genotype, plant traits were analyzed using full factorial linear models including all main effects and interactions (T × M × P). Type III ANOVA with sum-to-zero contrasts was used to accommodate the balanced design. Partial η² values were calculated to estimate the proportion of variance explained by each factor after accounting for interactions.

To characterize coordinated morphological changes, subsets of classic thermomorphogenic traits describing petiole morphology and rosette architecture were standardized and subjected to principal component analysis (PCA). We used a growth index defined as the first principal component (PC1) of standardized endpoint growth-related traits. Temperature-induced plasticity was quantified at the accession level as Δ(H–L), calculated from mean trait values at 28 °C (H) and 16 °C (L). Differences in plastic responses between warm- and cold-origin accessions were tested using two-sample *t*-tests. All plant statistical analyses were conducted in R (v4.5.1; [32]).

### Rhizosphere soil collection and sequencing

At the end of the temperature treatment, rhizosphere soil samples were harvested. Loosely attached soil was removed by gently shaking the roots. Roots were separated from shoots and transferred to 50 mL Falcon tubes containing 40 mL sterile deionized water. Tubes were gently inverted ten times to dislodge rhizosphere-associated soil. Roots were then removed, and the resulting wash solution was centrifuged at 4,000 × g for 10 min. The supernatant was discarded, and the pellet was resuspended in sterile water and transferred to 2 mL screw-cap tubes. After centrifugation at 20,000 rpm for 10 min, the supernatant was removed and the pellet was snap-frozen in liquid nitrogen and stored at −80 °C until analysis [17].

From the harvested rhizosphere material, three independent composite samples were generated for each treatment combination by randomly pooling three of the ten biological replicates. This resulted in 294 composite rhizosphere samples for downstream DNA extraction and amplicon sequencing. DNA was extracted using the DNeasy PowerLyzer PowerSoil Kit (QIAGEN, Hilden, Germany). DNA concentration was measured using a Qubit 2.0 fluorometer with the Qubit dsDNA BR Assay Kit (Thermo Fisher Scientific, MA, USA). Target regions were amplified using barcoded primers targeting the bacterial 16S rRNA V4 region and the fungal ITS2 region. Amplicons were sequenced on the Novogene Illumina platform to generate 250 bp paired-end reads. The statistics of sequencing data is shown in **Table S3**.

### Rhizosphere microbiome data processing and community analyses

Raw sequencing reads were processed using QIIME2 (v2025.7; [33]). Demultiplexed paired-end reads were merged using FLASH2 and denoised with DADA2 to generate amplicon sequence variants (ASVs; [34,35]). Chimeric sequences were removed using the uchime-denovo approach implemented in QIIME2. Taxonomic assignment was performed against SILVA v138.1 for bacterial 16S rRNA sequences and UNITE v9.0 for fungal ITS2 sequences. ASVs classified as chloroplast, mitochondria, or unassigned at the kingdom level were removed. To reduce sparsity, ASVs present in fewer than 10% of samples or with total abundance <20 reads across all samples were excluded. Unless otherwise stated, downstream analyses were performed on relative abundance tables normalized to unit total abundance. Bacterial and fungal communities were profiled with different amplicon markers and analyzed separately. We therefore limited comparisons between kingdoms to broad patterns such as community reorganization, microbiome-phenotype associations, and model performance, rather than direct contrasts in relative or absolute abundance.

#### Soil property associations with microbial community composition

Associations between soil physicochemical properties and major microbial phyla were assessed using Mantel tests. Relationships between microbial community composition and soil physicochemical variables were further examined using constrained ordination, with canonical correspondence analysis (CCA) applied to bacterial communities and redundancy analysis (RDA) applied to fungal communities. Mantel tests [36], CCA [37], and RDA [38] followed the linear constrained ordination framework implemented in the vegan package (v2.7-3) [39] in R.

#### Alpha and beta diversity analyses

Alpha diversity was calculated from ASV tables rarefied to 1,000 reads per sample using the Shannon index, Chao1, and observed richness. Beta diversity was quantified using Bray-Curtis dissimilarity based on relative abundance data, and principal coordinate analysis (PCoA) was used for ordination. Community composition was tested by PERMANOVA with 999 permutations, using temperature, microbial inoculum, plant genotype, and their interactions as explanatory terms. Multivariate dispersion was assessed separately by testing distances to group centroids with 999 permutations. Analyses were performed on the full dataset and within each temperature treatment (16 °C and 28 °C) in R using the vegan package (v2.7-3) [39–41].

#### Identification of recurrent temperature-responsive lineages

To identify microbial lineages consistently associated with temperature across experimental locations, ASV tables were aggregated at the family level. Within each sampling site, family-level relative abundances were compared between temperature treatments using linear models, and *P* values were adjusted for multiple testing using the Benjamini-Hochberg false discovery rate (FDR) procedure. Families were classified as recurrent temperature-responsive lineages when they showed a significant temperature effect (FDR < 0.05) in at least four independent locations with a consistent direction of response.

#### Microbial network construction and topological analysis

Microbial co-occurrence network analysis was used to examine whether elevated temperature was associated with changes in potential microbial association patterns in the rhizosphere. Microbial co-occurrence networks were constructed separately for the 16°C and 28°C treatments using the Molecular Ecological Network Analysis Pipeline (MENAP; Institute for Environmental Genomics, University of Oklahoma) [42]. Prior to network construction, taxa detected in fewer than 2% of samples were removed to reduce sparsity. Pairwise associations between relative abundances of different Phyla were calculated using Spearman rank correlations, and adjacency matrices were defined using the random matrix theory (RMT)-based threshold implemented in MENAP. Only correlations exceeding the RMT-derived cutoff were retained for network construction. Network topological properties, including node degree, clustering coefficient, modularity, transitivity, betweenness centrality, and eigenvector centralization, were calculated within MENAP to assess temperature-associated changes in network connectivity and compartmentalization. Modules were identified by modularity optimization. For downstream analyses, module eigengenes were defined as the first principal component of the standardized relative abundances of taxa within each module. Co-occurrence networks represent statistical associations; network differences were therefore interpreted as changes in potential microbial association structure rather than direct ecological interactions.

#### Quantification of changes in microbiome composition

Microbiome compositional change (Δβ) was quantified as Bray-Curtis dissimilarity relative to the corresponding home microbiome baseline. For each plant-microbiome combination, Δβ was calculated relative to the corresponding matched home rhizosphere community from the same temperature treatment at the end of the experiment, and was then classified by matching context as home or away. For accession-level analyses, Δβ values were averaged across biological replicates within each accession and matching context.

### Microbiome-phenotype associations and community-level coupling

Associations between accession-level growth index and microbiome compositional change (Δβ) were evaluated using Spearman rank correlations, conducted separately for home and away conditions. To assess robustness, leave-one-out sensitivity analyses were performed by iteratively excluding one accession at a time and recalculating the correlation coefficient, following a jackknife-type resampling approach [43]. Community-level microbiome-phenotype coupling was assessed using Mantel tests comparing Bray-Curtis dissimilarity matrices of bacterial or fungal communities with Euclidean distance matrices of multivariate plant phenotypes. Mantel tests were performed separately for home and away contexts.

### Predictive modeling framework

To quantify the association between microbiome composition and plant phenotypic variation, nested linear models were fitted using aggregated trait values (mean per T × M × P × matching combination). The baseline model included temperature (T), microbial inoculum (M), and plant genotype (P): *Trait* ∼ T + M + P

Extended models additionally included microbiome composition-derived predictors, namely bacterial and fungal compositional change (Δβ) and the first principal coordinate axis (PCoA1) derived from Bray-Curtis dissimilarities: *Trait* ∼ T + M + P + fungal Δβ + fungal PCoA1 + bacterial Δβ + bacterial PCoA1

To compare the relative contributions of the two microbial kingdoms, symmetric models were also fitted with either fungal or bacterial composition-derived predictors only. Model fit was evaluated using adjusted *R*² and Akaike’s Information Criterion (AIC), and nested models were compared using partial F-tests [44].

Predictive performance was further assessed by ten-fold cross-validation [45]. Cross-validated *R*² was calculated as: *R*²_CV = 1 − (Σ(*y*obs − *y*pred)² / Σ(*y*obs − ȳ)²) Semi-partial *R*², representing the unique variance explained by each focal predictor, was calculated from the change in residual sums of squares between full and reduced models [46]. All analyses were conducted in R (v4.5.1; [32]).

Throughout the manuscript and supplementary materials, sample IDs are given as Temperature_Microbial inoculum_Plant genotype. Temperature codes are L for 16 °C and H for 28 °C. Microbial inocula are coded as mSP1, mSP2, and mSP3 for the three Spanish microbial inocula, mSW1, mSW2, and mSW3 for the three Swedish microbial inocula, and mC for the sterilized-soil control. Plant genotypes are coded as pSP1, pSP2, and pSP3 for the three Spanish accessions, pSW1, pSW2, and pSW3 for the three Swedish accessions, and pC for *Col-0*. Official accession names corresponding to these soil collection sites and plant genotype codes are listed in **Table S1**.

### Soil covariate adjustment

#### Plant trait soil-covariate analysis

To test whether residual edaphic variation confounded inoculum-associated effects on plant traits, soil N and C/N ratio were summarized at the inoculum level and linked to the plant trait dataset by inoculum identity. For each trait, nested linear models were used to test the additional contribution of inoculum identity after accounting for temperature, plant genotype, soil N, and soil C/N ratio. The additional explanatory contribution of inoculum identity was quantified as partial R². In parallel, scaled multivariate plant traits were analyzed by partial redundancy analysis, with temperature, inoculum identity, and plant genotype as explanatory variables and soil N and C/N ratio included as conditioning variables. Term significance was assessed using 999 permutations.

#### Microbiome soil-covariate analysis

To assess whether residual soil N and C/N variation explained microbiome community structure, partial distance-based redundancy analysis was performed using Bray-Curtis dissimilarities for bacterial and fungal communities separately. Temperature, inoculum identity, and plant genotype were tested as explanatory variables while conditioning on soil N and C/N ratio. Conversely, the independent contribution of soil N and C/N ratio was tested after conditioning on temperature, inoculum identity, and plant genotype. Term significance was assessed using 999 permutations.

## RESULTS

### Temperature-responsive growth variation guides selection of climatically contrasting field sites

To guide field site selection for the factorial experiment, we conducted a two-step pilot screen using European *A. thaliana* accessions from the 1001 Genomes collection. First, 114 accessions from habitats spanning the warm and cold extremes of the species’ European climatic range were grown on medium plates at 16 °C and 28 °C to assess seedling thermoresponsiveness by analyzing the temperature-induced hypocotyl elongation, which is known to be a good proxy for various kinds of thermomorphogenic responses, also later in development [47]. This initial screen revealed substantial accession-level variation in relative hypocotyl elongation under warm versus cool ambient conditions (**Fig. 2a**). Based on these results, 48 accessions showing pronounced temperature-responsive growth differences, together with *Col-0* as a reference genotype, were selected for a second soil-based pilot experiment. This follow-up screen used the same cultivation conditions as the main experiment, with image-based phenotyping on day 11 of the temperature treatment of 28 °C compared to the 16 °C control treatment.

**Fig. 2.**
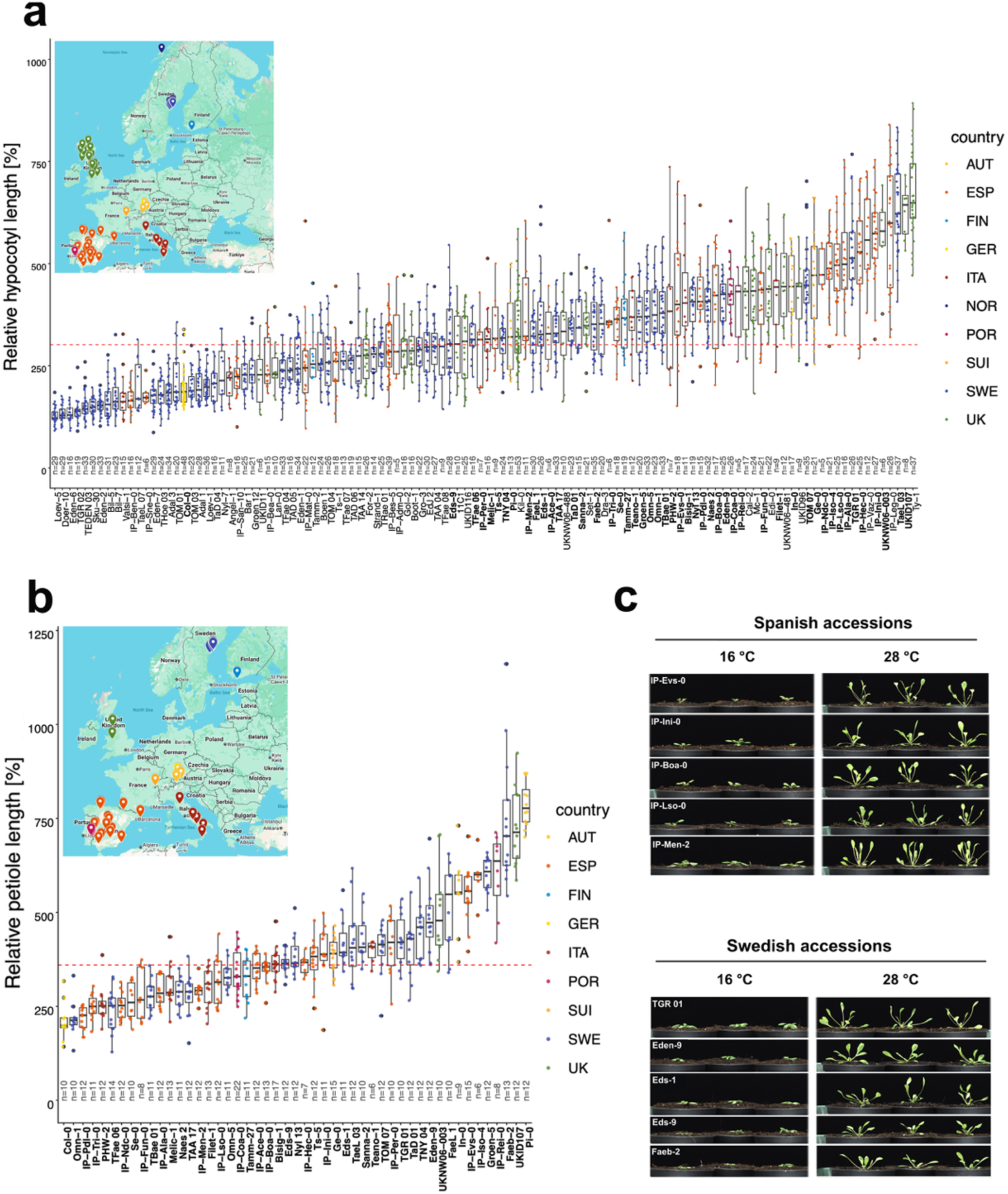
Two-step pilot screen of thermoresponsive growth used to guide field-site selection. (**a**) Initial screen of 114 European *A. thaliana* accessions from the 1001 Genomes collection grown on agar plates for 7 days to assess relative hypocotyl elongation under warm (28 ℃) versus cool ambient (16 ℃) conditions. Accessions in bold were used in (**b**): Follow-up soil-based screen of 48 accessions selected from the initial screen based on pronounced temperature-responsive growth differences, with *Col-0* (1001g ID: 6909) included as a reference genotype. Plants were grown under the same cultivation conditions as the main experiment and phenotyped by image analysis on day 11 of the temperature treatment to assess early vegetative responses, including relative petiole elongation. Accessions are ordered by their median relative response, and colors indicate country or region of origin. (**c**) Representative images of 10 accessions from the soil-based pilot screen, including five Spanish and five Swedish accessions, shown at day 11 of the temperature treatment under both 16 °C and 28 °C conditions. These images illustrate the range of early vegetative phenotypes observed among warm- and cold-origin materials under the two temperature treatments. The complete day-11 side-view images set for the 48 accessions is provided in **Fig. S2**.

It confirmed the broad variation in classic thermomorphogenic responses such as petiole elongation (**Figs. 2b, c and S2**), but revealed no clear correlation between temperature responsiveness and latitude. This is consistent with earlier work on the temperature dependence of flowering time, in which latitude alone proved a weak predictor of the underlying variation [48,49]. Accordingly, accession sampling was guided by the northern and southern range margins of *A. thaliana*-representing cold and warm climatic origins-while keeping major site characteristics broadly comparable.

We selected three field sites from central Spain and three from Sweden (**Table S1**) as sampling regions. The collected native accessions were then propagated for one generation in controlled greenhouse conditions to eliminate potential maternal effects caused by the native environments at the collection sites. Seeds of several individual plants per accession were bulked and used in the main experiment.

### Temperature strongly shapes vegetative plant growth and architecture

The soil-repopulation strategy was designed to reduce chemical differences among donor soils while retaining their microbial legacy. Before analyzing plant and rhizosphere responses, we first evaluated residual soil chemical variation in the repopulated substrate. Most physicochemical differences were substantially reduced after repopulation, although soil N content and C/N ratio remained distinguishable among inoculum origins (**Fig. S3**). These two variables were therefore included as covariates in subsequent sensitivity analyses.

We examined how temperature (T), microbial inoculum (M), and plant genotype (P) affected *A. thaliana* vegetative traits using a fully factorial P × M × T design (**Figs. 3 and S4-10**). To keep the main text focused, we highlight petiole length and plant area as representative univariate traits capturing architectural and size-related responses, respectively, and use multivariate phenotype axes to summarize coordinated trait variation. Detailed plots for the remaining measured traits are provided in the supplementary figures (**Figs. S4, S7-10**), and the corresponding Type III ANOVA results are summarized in **Table S4**.

**Fig. 3.**
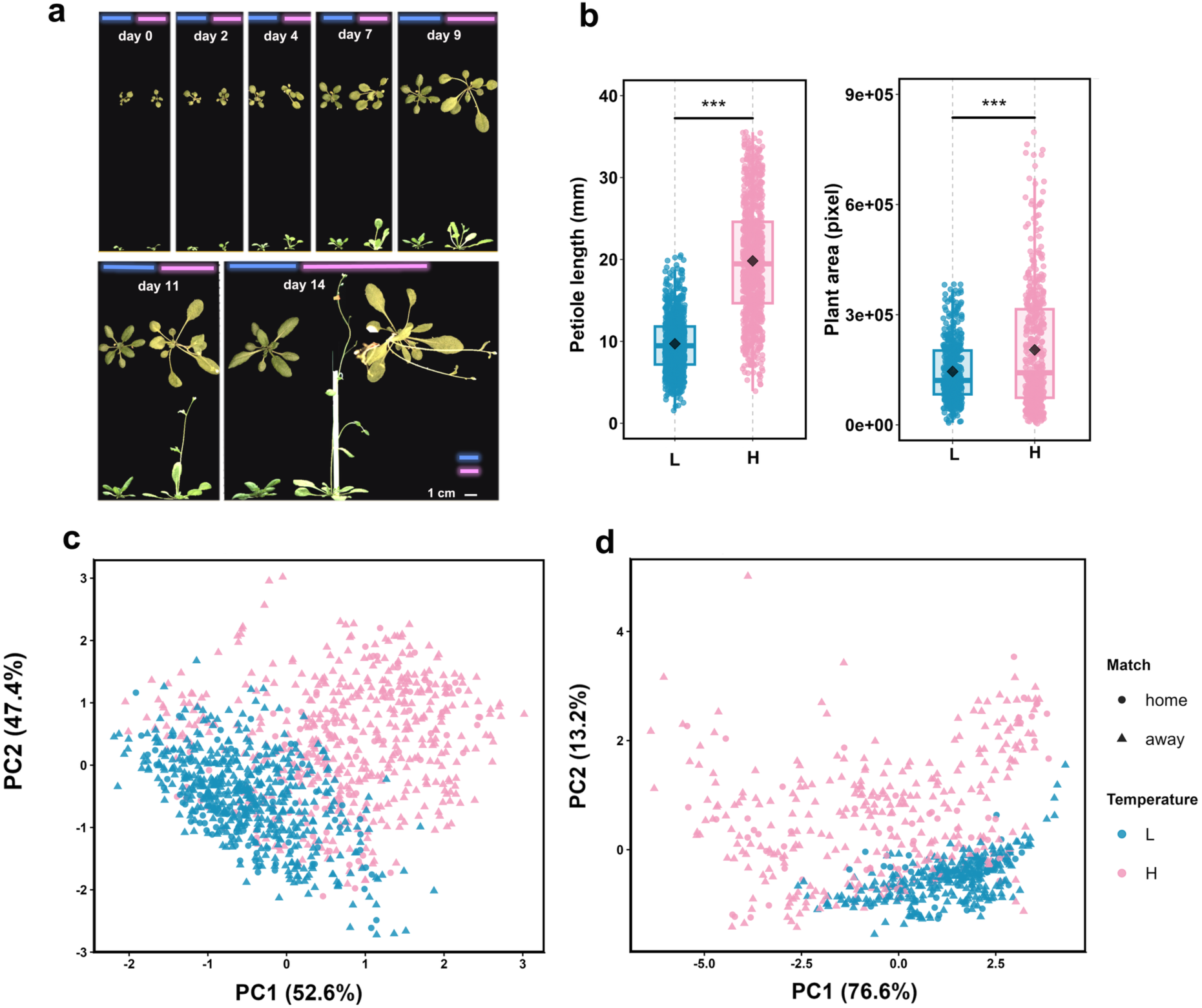
Plant growth and morphological traits in response to warming temperature. (**a)** Representative top-view and side-view images of wild-type *A. thaliana* accession *Col-0* (1001g ID: 6909) grown in control soil at 16°C (L) and 28°C (H). Representative images were taken at the indicated days (day 0, 2, 4, 7, 9, 11 and 14) after the start of the temperature treatment (day 0) to illustrate temperature-dependent differences in plant rosette architecture. (**b**) Petiole length and plant area measured at 14 days across the six wild accessions and *Col-0* under the two temperature treatments. Plant area was quantified using the LemnaTec phenotyping platform. Asterisks denote significance between temperatures after Benjamini-Hochberg correction (*FDR < 0.05, **FDR < 0.01, ***FDR < 0.001). (**c, d**) Principal component analysis (PCA) of petiole angle, petiole length traits (**c**), and rosette shape traits (**d**) across the six wild accessions and *Col-0* under the two temperature treatments. For host-microbiome matching analyses, home referred only to the original accession-soil pairing from the same field site of origin. Any non-native combination, including pairings within the same climatic region but from different sites, was classified as away. *Col-0* and the sterilized-soil control were excluded from home-away analyses. The plant trait analyses reported here were based on endpoint measurements taken after 14 days of temperature treatment.

Across the measured traits, temperature emerged as a major driver of plant morphology within the 14-day vegetative assay. At 28 °C, plants showed a coherent architectural shift characterized by petiole elongation and a more open rosette structure, which has been shown to enable evaporative cooling [50], relative to 16 °C (**Fig. 3a**). Univariate analyses confirmed strong temperature effects on both petiole length, plant area and other measured traits (**Fig. 3b and S4**). Plant genotype significantly affected plant phenotype, whereas microbial inoculum had comparatively weak main effects, with substantial overlap in trait distributions across inoculum treatments (**Figs. S5-S8**). Interaction analyses further showed that temperature responses varied among genotypes (T × P; **Figs. S5, S6 and S9**), but were broadly consistent across microbial inocula (T × M; **Figs. S5, S6 and S10**).

To capture coordinated effects on plant morphology, we performed principal component analysis (PCA) on petiole-related traits and rosette architecture. In both cases, the first principal component clearly separated plants grown at 16 °C and 28 °C (**Fig. 3c, d**), defining a dominant temperature-associated phenotypic axis. PC1 scores differed strongly between temperature treatments (**Fig. S11**), whereas differences attributable to home-away matching were comparatively modest at this stage, with substantial overlap in endpoint phenotype distributions across contexts. Variance partitioning further revealed a hierarchical structure of phenotypic control (**Fig. S12; Tables S4 and S5**). Across traits, plant genotype explained the largest proportion of total variance, followed by temperature and genotype-specific temperature responses (T × P). By contrast, microbial inoculum contributed relatively small independent effects.

Given the residual variation in soil N and C/N ratio, we further tested whether these edaphic covariates contributed to the weak inoculum-associated effects on plant traits. Soil-adjusted partial RDA showed that temperature, plant genotype, and inoculum identity all remained significant predictors of multivariate trait structure, but inoculum identity explained only a small fraction of the constrained variation relative to genotype and temperature (**Fig. S13**; **Table S6**). Trait-level nested models similarly showed that adding inoculum identity after temperature, plant genotype, soil N, and soil C/N explained only modest additional variation across individual traits, with partial R² values mostly below 0.02-0.03 (**Table S7**). Thus, residual soil N and C/N did not account for the weak but detectable inoculum-associated effects on plant traits.

We next asked whether climatic origin predicted stronger vegetative responses under elevated temperature. Responses were strongly genotype-dependent across all traits (**Fig. S14a, b**). Although most accessions showed increased trait values at 28 °C, the magnitude and direction of change varied substantially among genotypes. Warm-origin accessions did not consistently show greater vegetative growth than cold-origin accessions under elevated temperature. Cold-origin accessions exhibited larger temperature-induced trait shifts [Δ(H–L)] across traits (**Fig. S14c, d; Table S8**). One warm-origin accession (pSP2/IP-EVS-0_B) showed marked growth suppression at 28 °C; however, excluding this genotype did not alter the qualitative origin-dependent pattern of plastic responses (**Fig. S15; Table S9**).

### Warming reorganizes rhizosphere microbiome structure within soil- and plant-defined constraints

To determine how elevated temperature restructured rhizosphere microbial communities, we quantified the relative contributions of microbial inoculum (M), plant genotype (P), temperature (T), and their interactions to community composition. PERMANOVA revealed a hierarchical organization of rhizosphere microbiomes (**Fig. 4a, c**). Microbial inoculum explained the largest proportion of variation in both bacterial and fungal communities (52.25% and 55.10%, respectively), establishing inoculum origin as the dominant structural scaffold. Plant genotype contributed additional variation, particularly in fungal communities (2.78% for bacteria and 17.06% for fungi). Temperature accounted for a smaller but statistically significant fraction of variation in both kingdoms (1.05% and 1.40%, respectively).

**Fig. 4.**
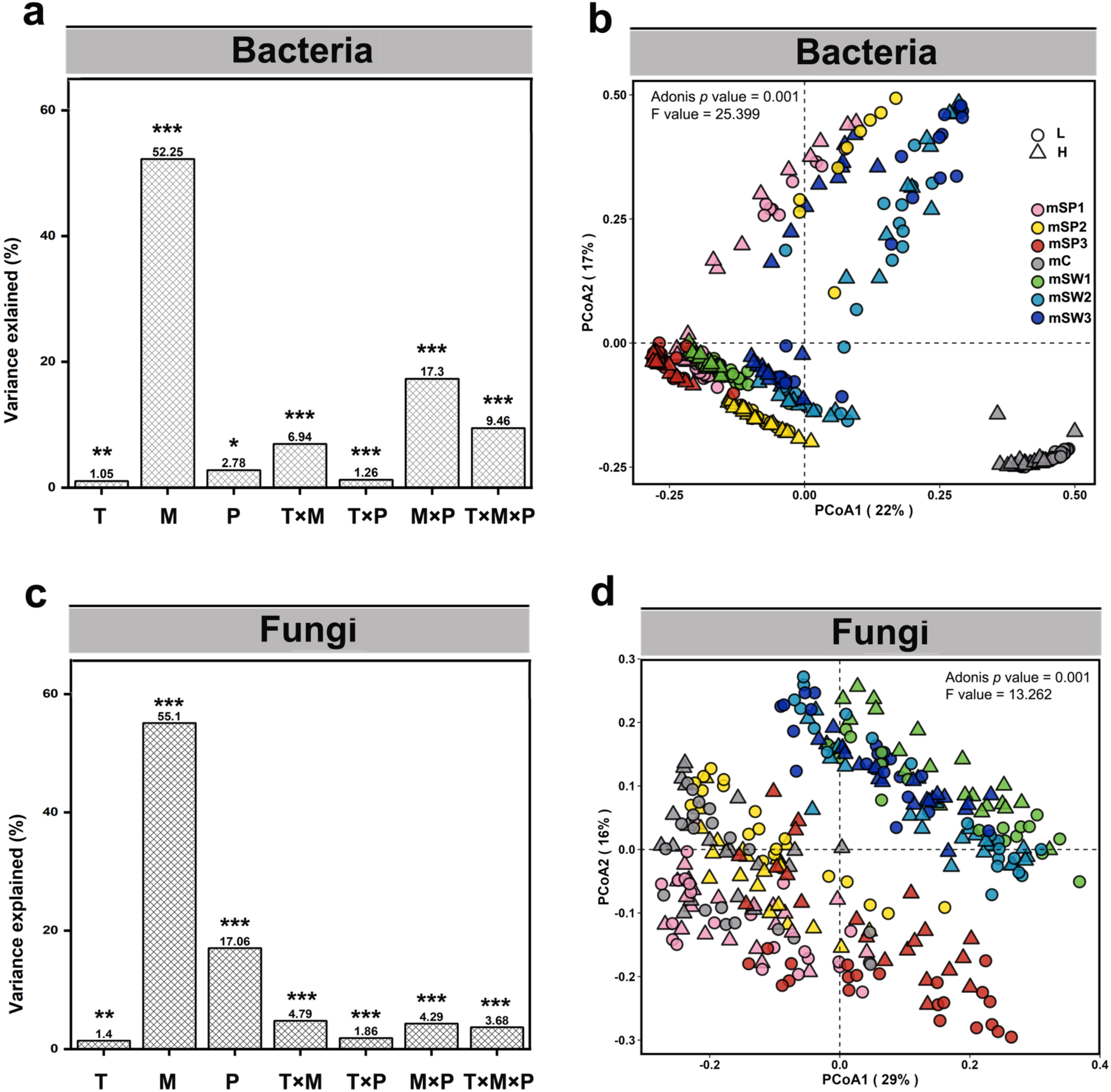
Effects of microbial inoculum, plant genotype, and temperature on rhizosphere microbiome composition. (**a, c**) Variance explained by microbial inoculum (M), plant genotype (P), temperature (T), and their interactions for bacterial (**a**) and fungal (**c**) community composition, estimated by PERMANOVA on Bray-Curtis dissimilarity matrices. Asterisks indicate term significance from permutation tests with 999 permutations (*FDR < 0.05, **FDR < 0.01, ***FDR < 0.001). (**b, d**) Principal coordinate analysis (PCoA) of bacterial (**b**) and fungal (**d**) rhizosphere communities based on Bray-Curtis dissimilarities. Points represent individual samples, colored by microbial inoculum origin and shaped by temperature treatment.

Principal coordinate analysis (PCoA) based on Bray-Curtis dissimilarity further illustrated these hierarchical patterns. When all samples were analyzed jointly, bacterial and fungal communities were separated primarily by microbial inoculum (**Fig. 4b, d**), with temperature not forming the dominant axis of variation. To resolve temperature-associated structure more explicitly, we performed temperature-stratified ordinations. Across inoculum-defined baseline configurations, elevated temperature induced reproducible directional shifts while preserving the dominant clustering by inoculum origin (**Fig. S16**). Comparable temperature-associated reorganization was also observed within plant genotype groupings, particularly in fungal communities (**Fig. S17**).

Multivariate dispersion analyses further showed that temperature altered within-group variability in both microbial kingdoms (**Fig. S18; Table S10**). Alpha diversity was structured primarily by inoculum origin and showed more limited, context-dependent responses to temperature and plant genotype (**Fig. S19**).

We next tested whether the residual variation in soil N and C/N ratio could account for inoculum-associated microbiome structure. After conditioning on these covariates, inoculum identity remained the dominant predictor of both bacterial and fungal community composition, whereas temperature and plant genotype explained smaller but significant components of variation (**Table S11**). Conversely, soil C/N explained no additional microbiome variation after conditioning on temperature, inoculum identity, and plant genotype, while soil N showed only a minor residual association with bacterial communities. These results indicate that the strong inoculum-associated structure of the rhizosphere microbiome was not reducible to residual variation in soil N or C/N.

### Recurrent temperature-responsive lineages and interaction reconfiguration under warming

Having established that warming induced reproducible compositional shifts within inoculum-and host-defined microbial baselines, we next asked whether this restructuring reflected stochastic turnover or the repeated selection of specific microbial lineages.

Temperature-responsive microbial families recurred across multiple geographic origins rather than being confined to individual sites (**Fig. S20; Table S12**). Several bacterial families, including *Pedosphaeracea*e, *Opitutaceae*, *Iamiaceae*, *Haliangiaceae*, *Vicinamibacteraceae*, *Microscillaceae*, *Gemmataceae*, and *Dongiaceae*, were repeatedly enriched under 28°C across locations. Fungal responses were more lineage-specific: *Nectriaceae* was recurrently enriched under 28°C, whereas *Didymellaceae*, *Cladosporiaceae*, *Pleosporaceae*, *Phanerochaetaceae*, and *Coniochaetaceae* were more frequently enriched under 16°C (**Fig. S20**).

To determine whether these lineage-level shifts were accompanied by changes in higher-order organization, we constructed molecular ecological networks separately for the 16 °C and 28 °C treatments (**Fig. 5a, b**). Both temperature regimes contained six major modules, but network topology differed substantially under warming. The 28°C network contained more nodes and links than the 16°C network and showed a higher average degree, indicating denser statistical associations among microbial taxa under elevated temperature (**Fig. 5c; Table S13**). The proportion of bacteria-fungi links increased under warming, whereas intra-kingdom links decreased. Topological indices further showed structural reconfiguration at elevated temperature (**Fig. S21**). Clustering coefficient, transitivity, and modularity were reduced at 28 °C, whereas overall connectedness increased. The proportion of negative links also increased under warming, accompanied by a decrease in positive links (**Fig. 5c**). Network properties under both temperature regimes differed from those of randomized networks, indicating non-random organization.

**Fig. 5.**
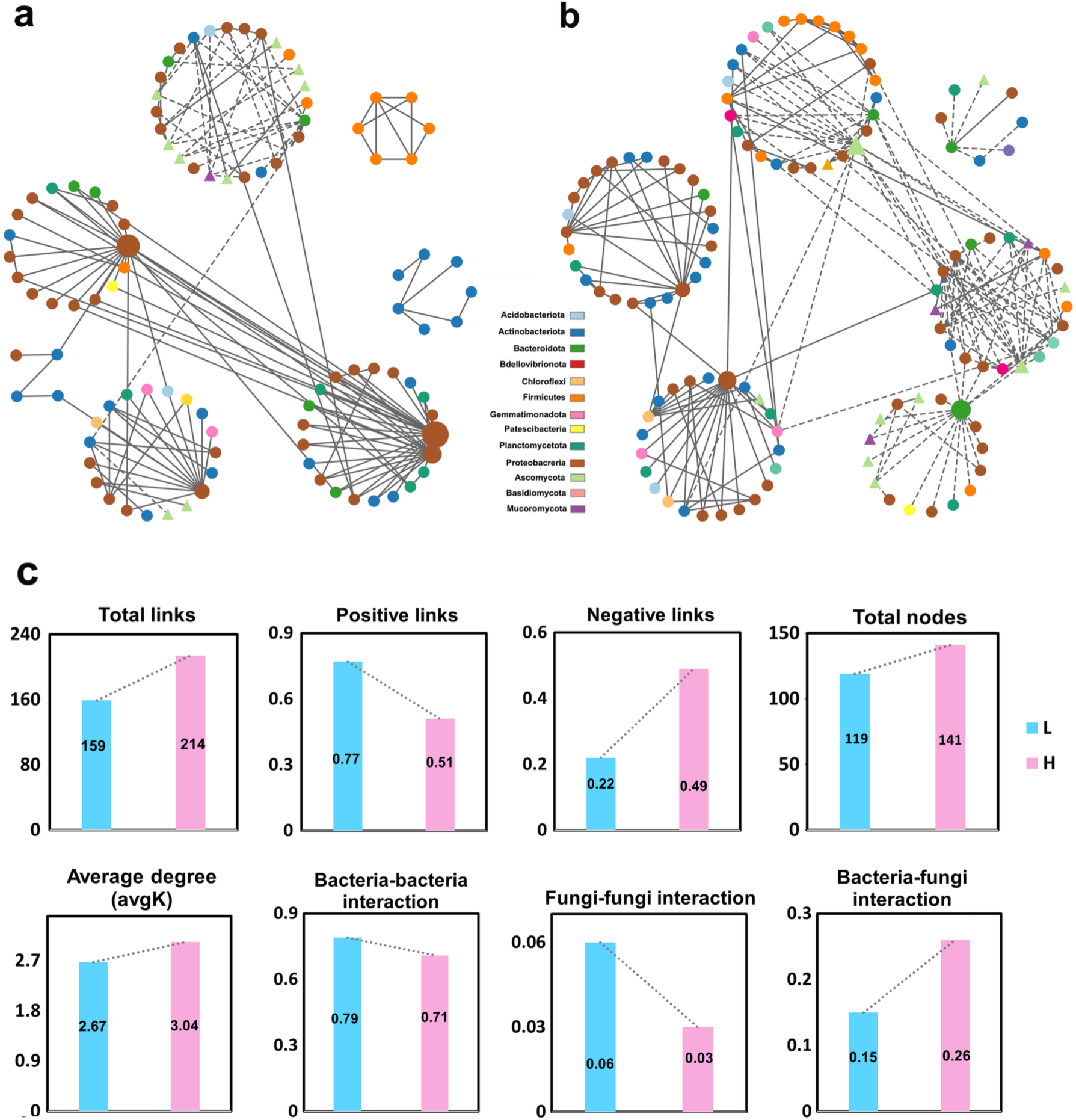
Warming reconfigures rhizosphere microbial co-occurrence network structure. (**a, b**) Temperature-specific co-occurrence networks constructed for rhizosphere microbial communities at 16 °C (**a**) and 28 °C (**b**) using the Molecular Ecological Network Analysis Pipeline (MENAP). Nodes represent bacterial or fungal taxa, and edges represent significant Spearman correlations retained after application of the random matrix theory threshold. Node colors indicate taxonomic groups. (**c**) Summary of selected network properties for the 16 °C and 28 °C networks, including the numbers of nodes and links, average degree, and the proportions of positive versus negative links and intra- versus inter-kingdom links. Additional topological indices are shown in **Fig. S21**, and full network statistics are provided in **Table S15**.

Despite these compositional and network-level shifts, predicted functional profiles remained broadly similar between temperature treatments. PICRUSt2-inferred bacterial metabolic pathways showed substantial overlap between 16 °C and 28 °C communities (**Fig. S22a**), and FUNGuild-based trophic assignments revealed comparable guild composition across temperatures (**Fig. S22b**).

### Microbiome structural stability associates with plant performance in a context-dependent manner

To assess whether host-microbiome matching contributes to rhizosphere community stability, we quantified microbial compositional change (Δβ) under home and away conditions across temperature treatments. Lower Δβ values indicate greater compositional stability (**Fig. S23**). Bacterial communities showed a significant home-away difference at 16°C, with higher compositional turnover under home than under away conditions. This direction did not support a simple home-stability pattern. The home-away difference disappeared at 28°C and was not observed in fungal communities, indicating that matching effects were weak, kingdom-specific, and temperature-dependent.

We next examined whether microbiome compositional change (Δβ) covaried with accession-level growth index under warming. At the accession level, Δβ was not significantly associated with growth index under either home or away conditions (**Fig. 6a, b**). The home subset showed a directionally consistent positive trend, whereas the away subset showed no clear directional pattern. Sensitivity analyses excluding individual genotypes indicated that the home trend remained directionally consistent, although effect sizes varied across iterations (**Table S14**).

**Fig. 6.**
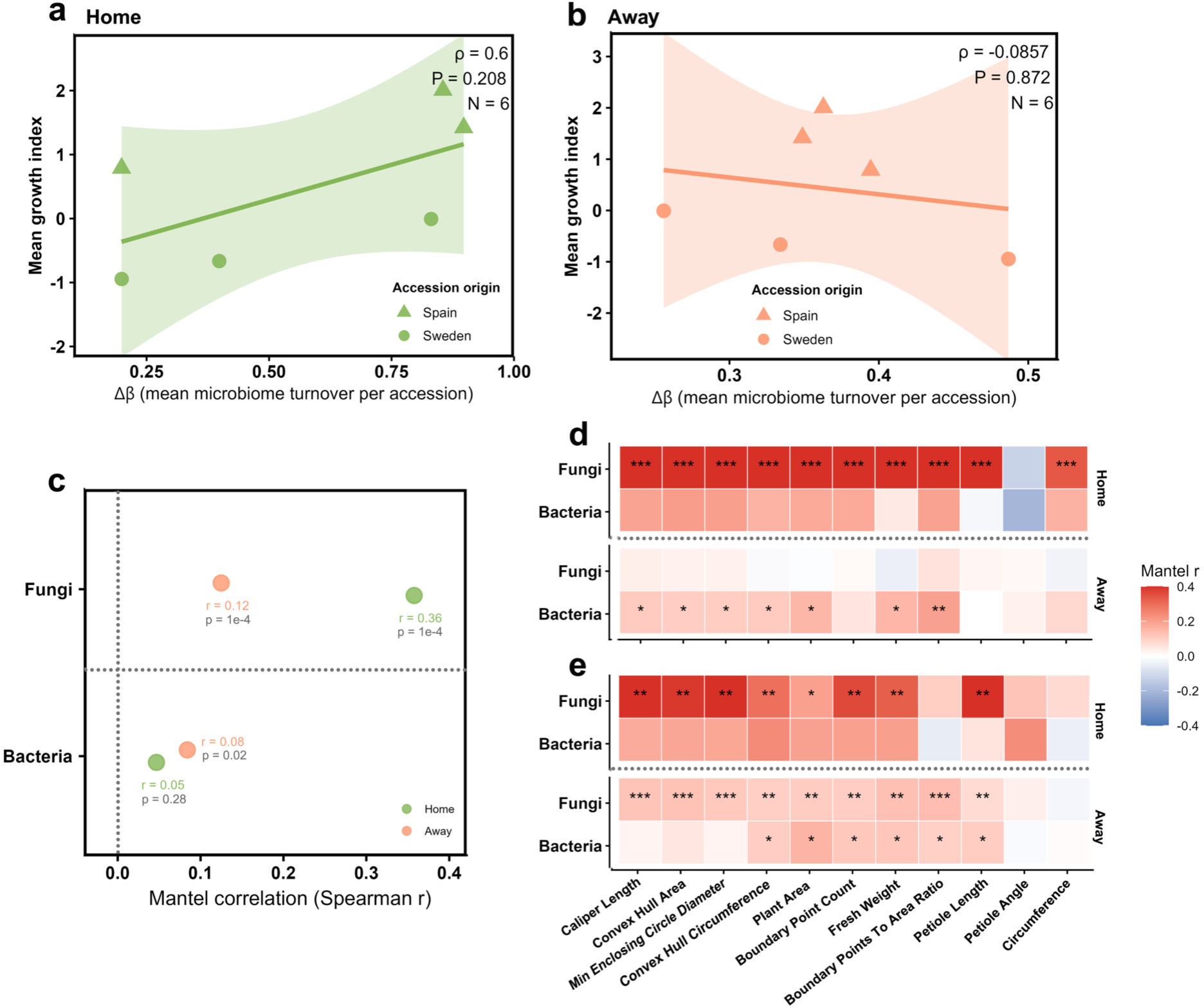
Microbiome-phenotype associations across temperature and matching contexts. (**a, b**) Relationships between accession-level plant growth index and rhizosphere microbiome compositional change (Δβ) under home (**a**) and away (**b**) conditions. The growth index was defined as the first principal component (PC1) of plant growth traits. Each point represents one accession mean. Lines indicate fitted linear trends; Spearman’s correlation coefficients and associated *P* values are shown in the panels. *Col-0* and the sterilized-soil control were excluded from home-away analyses. (**c**) Mantel correlations between microbiome Bray-Curtis dissimilarity and multivariate plant phenotypic dissimilarity for bacterial and fungal communities under home and away conditions. Points indicate Mantel’s *r*; colors denote matching context. The dashed vertical line indicates *r* = 0. (**d, e**) Trait-resolved Mantel heatmaps showing correlations between microbiome dissimilarity and individual plant traits (see also in **Table S2**) at 16 °C (**d**) and 28 °C (**e**). The plant trait analyses reported here were based on endpoint measurements taken after 14 days of temperature treatment. Tile color represents Mantel’s *r*. Asterisks indicate significance after Benjamini-Hochberg correction within each dataset × context × temperature stratum (*FDR < 0.05, **FDR < 0.01, ***FDR < 0.001).

Community-level analyses further revealed kingdom-specific differences in microbiome-phenotype coupling. Mantel tests showed significant associations between fungal community dissimilarity and multivariate plant phenotypic dissimilarity under both home (*r* = 0.358, *P* = 1 × 10⁻⁴) and away (*r* = 0.125, *P* = 1 × 10⁻⁴) conditions (**Fig. 6c**). In contrast, bacterial community dissimilarity showed weaker associations, which were not significant under home conditions (*r* = 0.0468, *P* = 0.267) and were modest under away conditions (*r* = 0.0840, *P* = 0.0218). Trait-resolved Mantel analyses yielded similar patterns, with stronger and more consistent fungal coupling across temperature and matching contexts (**Fig. 6d, e**).

To connect interaction architecture with phenotypic variation, we incorporated network module eigengenes into regression models (**Fig. S24**; **Table S15**). Among all tested modules, only L_M8, derived from the 16°C network, was retained as a significant module-level predictor after controlling for environmental and host variables. L_M8 was significantly associated with plant area, phenotype PC1, and petiole length, and was also related to the fungal structural axis PCoA1 (**Fig. S24**). No module-level predictors were detected at 28°C.

### Fungal community composition improves prediction of short-term vegetative performance under warming

Building on the inoculum- and host-constrained reorganization of rhizosphere microbial communities and the context-dependent microbiome-phenotype relationships described above, we next examined whether the composition of the fungal rhizosphere microbiome at the end of the experiment improved the prediction of plant phenotypic variation.

Linear models revealed significant associations between fungal PCoA1 and plant area, phenotype PC1, and petiole length (**Fig. 7a-c**). In each case, endpoint trait values varied systematically along fungal community composition PCoA1. By contrast, equivalent models using bacterial community composition did not show consistent or significant associations (**Table S16**). We next quantified model improvement after adding microbiome composition-derived predictors to the baseline environmental model (T + M + P). Inclusion of bacterial and fungal composition-derived metrics (Δβ and PCoA1) improved model fit across traits, with the strongest improvement observed for plant area (ΔAdjusted R² = +0.100; ΔAIC = −18.64; **Fig. S25**; **Table S17**). Improvements were more modest for phenotype PC1 (ΔAdjusted R² = +0.029; ΔAIC = −3.37) and petiole length (ΔAdjusted R² = +0.017; ΔAIC = −3.35; **Table S17**). Inclusion of host–microbiome origin matching did not further improve model performance. To compare microbial kingdoms, we fitted symmetric models incorporating either fungal or bacterial composition-derived predictors. These kingdom-specific models showed that fungal predictors generally provided stronger model improvement than bacterial predictors, particularly for plant area and phenotype PC1 (**Table S16**). Across traits, adding fungal composition-derived predictors (fungal Δβ and fungal PCoA1) to the baseline model consistently improved model fit, whereas adding bacterial composition-derived predictors (bacterial Δβ and bacterial PCoA1) produced minimal or inconsistent gains. This contrast was most pronounced for plant area, for which fungal predictors increased adjusted R² and reduced AIC, while bacterial predictors did not improve the model (**Table S16**).

**Fig. 7.**
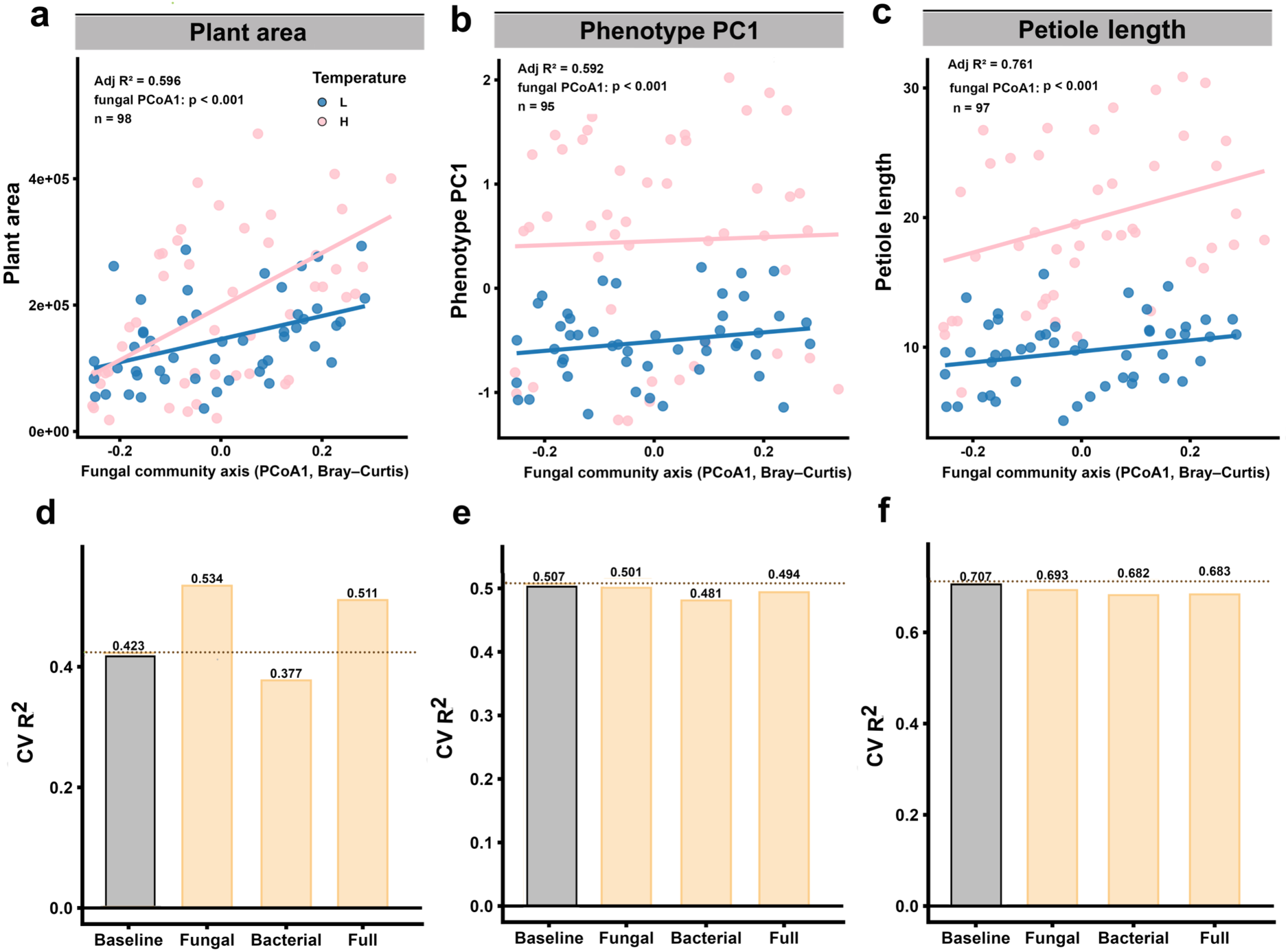
Fungal composition-derived predictors more consistently predict endpoint plant phenotypes than bacterial predictors. (**a-c**) Relationships between the fungal community composition (PCoA1 derived from Bray-Curtis dissimilarity) and plant area (**a**), phenotype PC1 (**b**), and petiole length (**c**). Each point represents the mean of one treatment combination. Colors indicate temperature treatment. Solid lines show fitted linear relationships across all observations. Statistical annotations report the effect of fungal PCoA1 in full linear models including temperature (T), microbial inoculum (M), plant genotype (P), host-microbiome matching status, and bacterial composition-derived predictors. (**d-f**) Ten-fold cross-validated predictive performance (*R*² 𝐶𝑉) of nested linear models for plant area (**d**), phenotype PC1 (**e**), and petiole length (**f**). The baseline model included T, M, and P. Extended models additionally included fungal composition-derived predictors (Δβ and fungal PCoA1), bacterial composition-derived predictors (Δβ and bacterial PCoA1), or both. Bars show cross-validated *R*² values, with exact values indicated above bars.

Cross-validation analyses further showed that adding fungal community composition axes produced reproducible predictive gains across traits (**Fig. 7d-f**), with consistent increases in cross-validated R² relative to the baseline model and to models adding bacterial community composition axes. Observed-versus-predicted relationships showed good correspondence between fitted and measured values (**Fig. S26**), supporting the robustness of fungal composition-derived predictors. In full models including both fungal and bacterial axes (**Table S16**), fungal PCoA1 remained a significant predictor for all three traits, whereas bacterial PCoA1 was not significant. Semi-partial R² analyses and standardized regression coefficients (**Fig. S27; Table S18**) likewise indicated larger unique and standardized effects for fungal than bacterial composition-derived predictors.

## DISCUSSION

In reciprocal plant-soil systems, an important unresolved question is whether warming disrupts historically structured plant-microbiome associations or instead reshapes them within existing ecological baselines [3,22,51]. In this study, warming did not simply override pre-existing plant–soil–microbiome relationships, but reshaped them in ways that remained dependent on the biological structure already present in the system. Temperature effects were clearly detectable across both plant phenotypes and rhizosphere communities, yet these effects were expressed through persistent host- and soil-associated baselines rather than as uniform responses. Our results also provided only limited support for simple origin-based matching of plant genotypes and inocula as a general explanation for plant performance. Instead, microbiome composition, particularly fungal community composition, was more closely associated with endpoint vegetative performance than host-microbiome origin matching alone. Therefore, plant responses to warming in this system were best understood as emerging from thermal reorganization acting on pre-existing biological structure, rather than from temperature effects in isolation.

### Temperature strongly reprograms plant phenotype while reorganizing rhizosphere communities within established ecological scaffolds

Elevated temperature was a major driver of both plant phenotype and rhizosphere community structure, but in neither case did it erase the pre-existing biological context in which these responses were expressed. On the plant side, warming induced a coherent shift in architecture, including petiole elongation and coordinated changes in rosette-related traits, consistent with well-described thermomorphogenic responses in *A. thaliana* [6,47]. Plant genotype explained a larger share of endpoint vegetative trait variation than microbial inoculum. At 28°C, cold-origin accessions tended to produce larger final rosettes than warm-origin accessions, indicating that warm-origin accessions did not show a general endpoint phenotype advantage under elevated temperature (**Figs. 3 and S14**). Under our experimental conditions, these patterns provide limited support for a simple local climatic adaptation interpretation [16,17,52,53]. Instead, the results show strong genotype-specific thermal responses, but provide limited support for an adaptation to local climate [47,54]. One caveat is of course implemented by the experimental design with an unusually high ambient temperature at both the northern and southern sites during the monitored developmental stages. However, all phenotypes reported here were measured within 14 days of the temperature shift, a standard window for *Arabidopsis* thermomorphogenesis assays, under conditions broadly applied in the thermomorphogenesis literature. These inferences are largely limited to vegetative developmental stages under constant long-day photoperiod and two setpoint temperatures; later developmental stages when microbial effects may potentially become more pronounced and fluctuating regimes were not assessed [55,56].

A comparable pattern was evident in the rhizosphere microbiome. Although temperature significantly shifted the composition of both bacterial and fungal communities, these shifts were secondary to the strong structuring effect of the origin of the microbial inoculum (**Figs. 4, S16 and S17**). In both kingdoms, soil-derived inoculum explained most of the variation in community composition, whereas temperature imposed reproducible directional changes within this inoculum-defined framework. This pattern is consistent with previous work showing that environmental filtering can reshape plant-associated microbiomes without fully overriding the influence of source community composition, edaphic history, or geographically structured microbial pools [16,17,57,58]. Warming therefore acted less as a replacement force than as a modifier of an already structured assembly background.

Plant and microbial responses to warming were parallel in one important respect: both were strongly influenced by pre-existing variation. Plant phenotypic responses depended on accession identity, whereas microbial responses primarily depended on inoculum background, with additional host effects that were particularly evident in fungi (**Figs. 3 and 4**). Elevated temperature did not impose a uniform response across plant-soil systems, but instead interacted with historically and biologically structured baselines to produce context-dependent outcomes. This view is consistent with broader work showing that responses to climate-related change are filtered through pre-existing host and community structure rather than determined by temperature alone [17,58,59]. It also helps frame the downstream links between microbiome structure and plant performance, which in this system emerged against persistent host and soil constraints.

### Warming selected recurrent microbial lineages and reorganized rhizosphere community structure

The microbial response to warming was clearly directional rather than random. Across sampling sites, the abundance of several bacterial and fungal families repeatedly differed between temperature treatments, often with consistent response directions (**Fig. S20**). Such deterministic filtering under elevated temperature, indicate that warming did not simply increase compositional noise but repeatedly favored or disfavored particular lineages across distinct plant-inoculum contexts [60–63]. Thermal change may therefore constrain rhizosphere assembly by selecting a narrower subset of community configurations able to persist under warmer conditions [42,64].

This selective restructuring was also evident at the level of community organization. Under 28 °C, the rhizosphere network showed higher connectivity, a greater proportion of bacteria-fungi links, and lower clustering, transitivity, and modularity than at 16 °C (**Figs. 5 and S21**). Although co-occurrence networks do not identify the causal ecological interactions, these shifts indicate that warming was associated with a less compartmentalized and more reconfigured community structure [42]. Elevated temperature therefore changed not only which taxa were favored, but also how bacterial and fungal groups were arranged relative to one another within the rhizosphere. In biological terms, this points to reorganization at the level of community architecture, with potential consequences for how different microbial groups respond jointly to the rhizosphere environment under warming.

Notably, these taxonomic and structural changes were not accompanied by strong divergence in predicted functional profiles (**Fig. S22**). Warming therefore appeared to reshape community membership and organization more strongly than broad predicted functional categories, a pattern consistent with the partial decoupling of taxonomic and functional responses often observed in microbial communities [63,65,66]. Given the limitations of PICRUSt2- and FUNGuild-based inference, this result should be interpreted cautiously [67,68]. Repeated lineage-level turnover, network rewiring, and relatively stable predicted functional structure point instead to structural reassembly within a constrained functional space rather than wholesale functional replacement [63,65]. This matters for the interpretation of later microbiome-phenotype relationships, because plant performance under warming was associated with a microbiome that had already been selectively filtered and reorganized, rather than simply displaced.

### Host-microbiome matching was limited and temperature-dependent

Previous work has suggested that long-term associations between plants and their local soil microbiota can generate home-field advantage or contribute to microbe-mediated local adaptation, potentially improving host performance or microbiome persistence in native combinations [16,69,70]. Empirical support for such effects, however, has often proved to be context-dependent rather than universal [19,71,72]. In our system, home-away effects on rhizosphere turnover were limited, being detectable only in bacterial communities at 16 °C and disappearing under warming, whereas fungal communities showed no corresponding turnover signal (**Fig. S23**). Rather than supporting a general home-associated stabilization of the microbiome, these results indicate that matching effects on community composition were weak, temperature-dependent, and kingdom-specific. Consistent with this, host-microbiome origin matching did not improve the prediction of endpoint plant phenotypes in our models. Under lower temperature, historically associated host-microbial inoculum combinations may retain enough ecological continuity for differences in bacterial turnover to emerge, whereas stronger thermal filtering under warming may reduce the expression of these finer-scale association [16,17].

### Microbiome-phenotype relationships were better captured by community structure than by matching status

If identity matching was not a general explanation for rhizosphere stability, it was also unlikely to be the main route through which the microbiome influenced plant phenotype. Instead, our results indicate that microbiome-phenotype coupling was captured more effectively by variation in community structure than by the simple distinction between home and away combinations. This was already apparent in the Mantel analyses, where fungal community dissimilarity showed stronger and more consistent associations with plant phenotypic dissimilarity than bacterial community dissimilarity under both matching contexts (**Fig. 6c-e**). In this system, the effect of the microbiome on plant phenotype therefore depended less on whether hosts and microbiota shared a common origin and more on how rhizosphere communities were compositionally organized [58,73–75].

Across analyses, fungal communities showed stronger plant-related structuring during assembly (**Fig. 4a, c**), stronger community-level coupling with plant phenotypic variation in the Mantel tests (**Fig. 6c-e**), and more consistent predictive gains when fungal composition-derived variables were added to nested linear models and cross-validation analyses (**Figs. 6 and S25-S27**; **Tables S16-S18**). By comparison, bacterial composition-derived predictors contributed more weakly and less consistently. This contrast suggests not simply that fungi were “more important,” but that fungal community organization may have integrated host-responsive and environmentally filtered variation in a way that remained more directly linked to plant growth outcomes under the conditions tested. As fungi often influence plants through traits such as nutrient acquisition, carbon exchange, and close physical association with roots, shifts in fungal community configuration may more readily translate into differences in plant growth than equivalent compositional shifts in bacteria [25,58,73–75].

The temperature dependence of this pattern further clarifies the ecological scale at which microbiome effects were expressed. At 16°C, the only significant module-level predictor, L_M8, was associated with multiple plant traits and was also linked to the fungal community composition (**Fig. S24**), suggesting that phenotype-relevant microbiome effects were expressed through more localized interaction structure under cooler conditions. Under warming, however, the most robust and repeatable predictive signal came from the broader fungal community composition rather than from individual network modules (**Figs. 7 and S24**). This shift suggests that warming altered not only microbiome composition, but also the organizational level at which microbiome variation affected plant phenotype and growth. Rather than uniformly amplifying or suppressing microbiome effects, warming appears to have shifted the relative importance of local interaction features of the microbiome toward emergent properties of the fungal community as a whole [42,76]. Therefore, the key result is not simple origin matching, but rather that plant phenotypes under warming were more closely associated with the overall organization of the rhizosphere microbiome, especially its fungal component [77–79].

### Implications for plant-microbiome coupling under warming

These patterns suggest that, under warming, rhizosphere organization may provide a more useful framework than host-microbiome origin matching for understanding how microbiomes relate to plant performance. In particular, the consistent signal associated with fungal community structure points to a level of organization that may be informative for future efforts to identify microbiome features linked to plant function under changing temperature conditions [69,78]. Rather than focusing only on whether native host-microbiome combinations are maintained, an important next step will be to determine which community states remain associated with plant performance as environments shift.

Future work should test these associations more directly and in ways relevant to microbiome-based improvement of plant performance. In particular, cross-inoculation experiments with reduced-complexity or synthetic communities could help determine whether the fungal composition-derived signal identified here reflects a causal contribution to plant growth under warming or a broader rhizosphere state associated with improved performance [80,81]. Direct functional measurements and validation under field conditions or fluctuating thermal regimes will also be needed to assess how stable and transferable these structure-based relationships are outside controlled environments.

## CONCLUSIONS

Under our controlled growth chamber conditions, elevated temperature was associated with reproducible shifts in plant architecture and rhizosphere community structure within host- and inoculum-determined baselines. Home-away matching explained little of the observed phenotype variation, whereas broader microbiome organization, particularly fungal community composition, carried more consistent phenotype-related signal under warming. Our results therefore indicate that plant responses to elevated temperature were linked less to origin-based matching than to the organization of the rhizosphere microbiome under changing thermal conditions. These findings underscore the value of considering community organization, rather than origin alone, in studies of climate-sensitive plant-microbiome relationships.

## DECLARATIONS

### Availability of data and material

The amplicon sequencing data generated during the current study are available at the National Center for Biotechnology Information (NCBI) in the Sequence Read Archive under the BioProject ID: PRJNA1470703. The sequencing data have been uploaded to NCBI and can be accessed via http://www.ncbi.nlm.nih.gov/bioproject/1470703.

### Authors’ contributions

MQ, MT, KK, CR, and IG conceived the project. XL, JT, SH, and MQ designed the experiments. XL and JT performed the experiments. LEL established the crop score imaging and image analysis. CAB, JÅ, JSP, and MQ collected plant and soil samples. AB, XL, and JT analyzed the soil samples. XL, JT and JZ analyzed the data. XL wrote the manuscript. MQ and JZ supervised and revised the manuscript. All authors read and approved the final manuscript.

## ACKNOWLEDGEMENTS

We thank Raphaela Pilz for her diligence in establishing the plant phenotyping system, and Robert Mikutta and Klaus Kaiser for supervising the soil analysis in their lab.

## SUPPLEMENTARY INFORMATION

Supplemental material is available online only.

Additional File 1, PDF: Supplemental Figures S1 to S27.

Additional File 2, Excel file: Supplemental Tables S1 to S18.

