## Supplemental Figures for "Fungal community composition links rhizosphere microbiome organization to plant phenotype in response to moderate warming"

Running title: *Arabidopsis* phenotype and rhizosphere microbiome under warming

**This Supplemental file includes:**

Figures S1 to S27

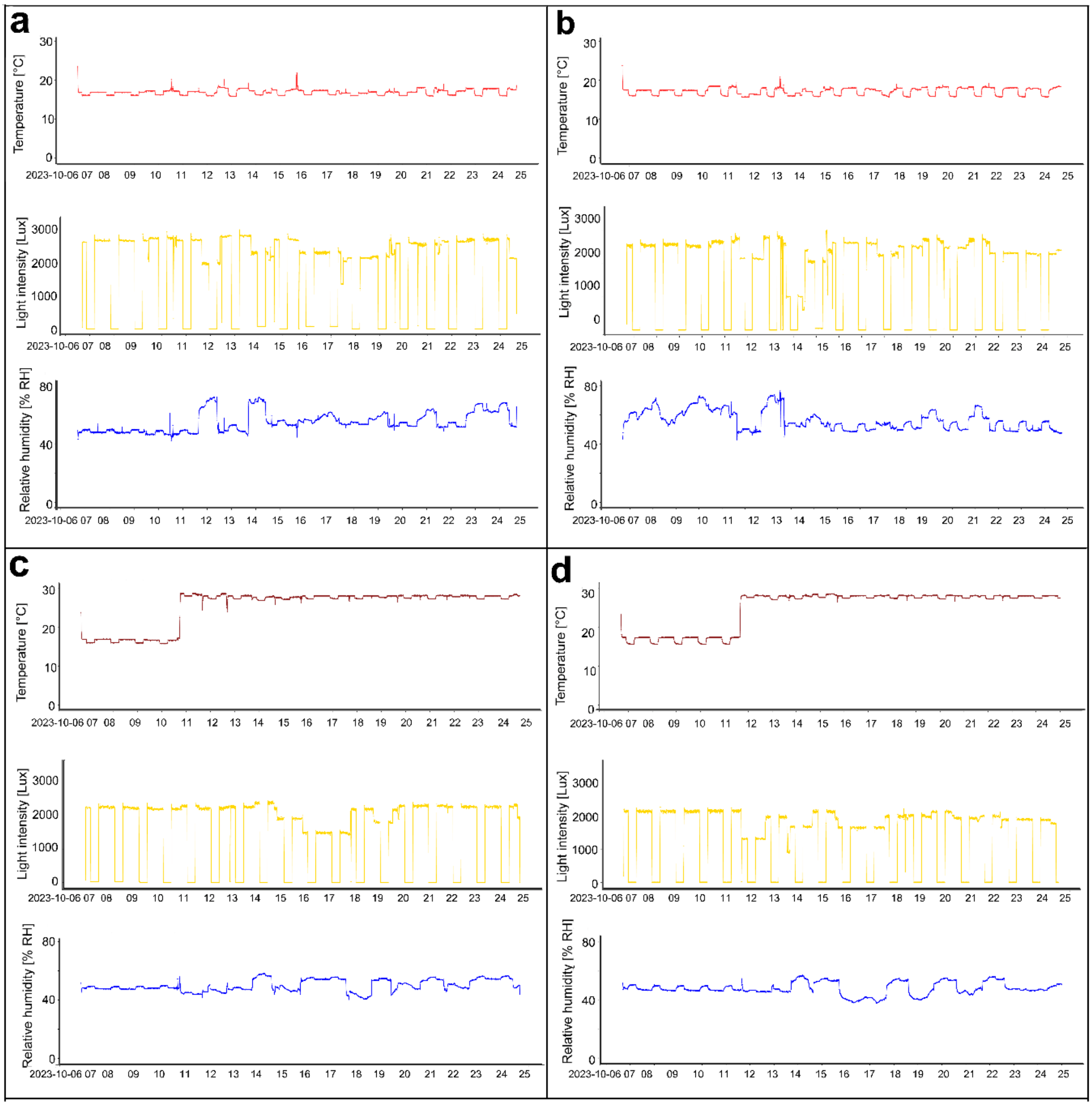


**Fig. S1. Logged environmental conditions in growth chambers.** Environmental parameters were continuously recorded to verify chamber comparability during the experiment. Panels **a** and **b** show the two chambers assigned to the low-temperature treatment (16 °C), whereas panels **c** and **d** show the two chambers assigned to the high-temperature treatment (28 °C). Logged parameters included relative humidity, photosynthetically active radiation (PAR), light spectrum, and air temperature. These records were used to confirm that replicate chambers within each temperature treatment maintained comparable environmental conditions.

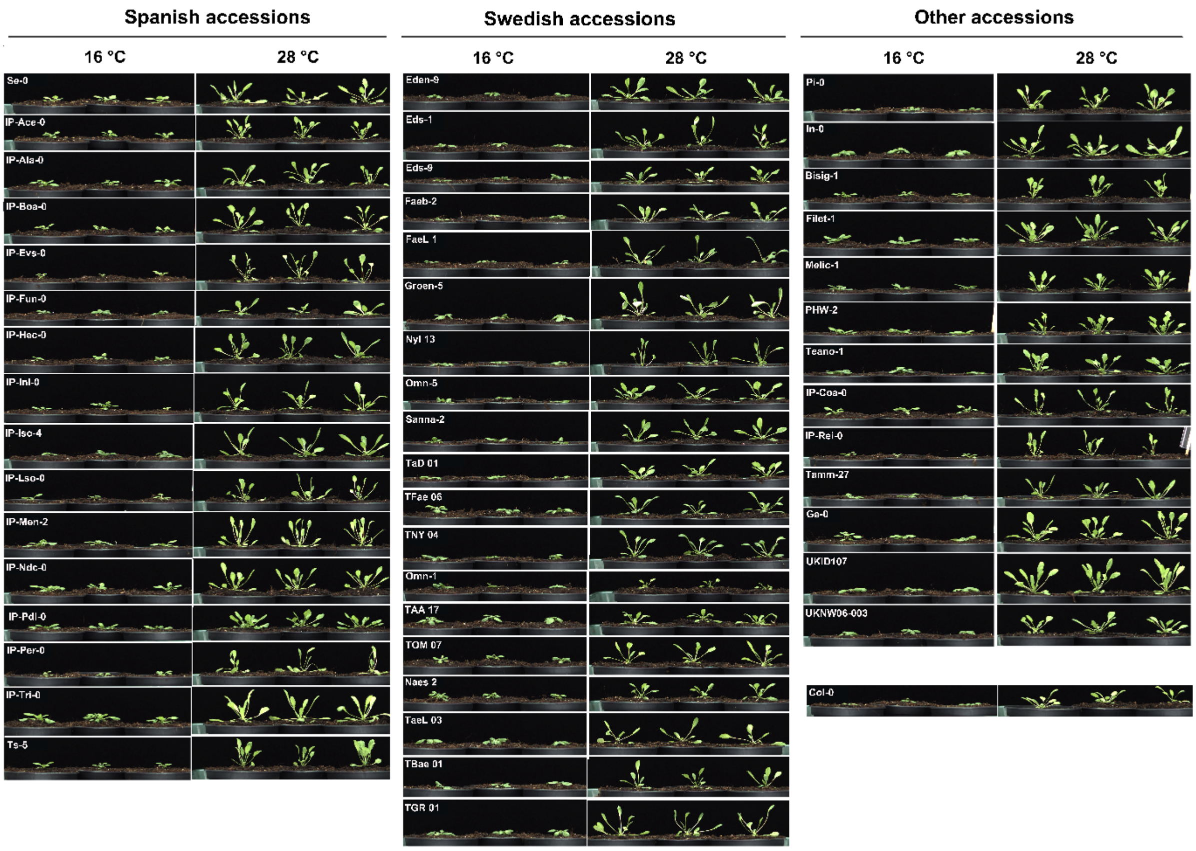


**Fig. S2. Day-11 side-view images of all accessions included in the soil-based pilot screen.** Side-view images show the 48 *A. thaliana* accessions selected from the initial agar-plate screen for the follow-up soil-based pilot experiment, with *Col-0* (1001g ID: 6909) included as a reference genotype. For each accession, plants grown at 16 °C and 28 °C are shown at day 11 of the temperature treatment. The soil-based screen used the same cultivation conditions as the main experiment and was used to visualize accession-level variation in early vegetative architecture and warm-versus-cool growth responses. This complete image panel complements the quantitative trait summary shown in **Fig. 2b**.

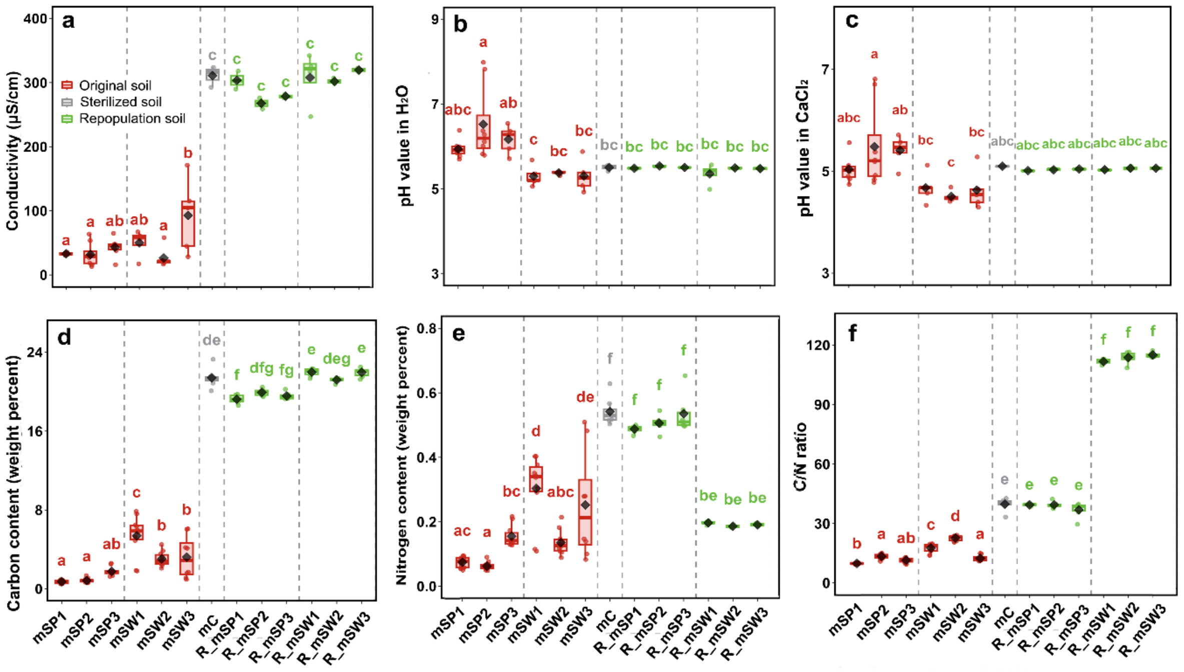


**Fig. S3. Soil physicochemical properties of original donor soils and repopulated soils and their associations with microbial community structure.** Electrical conductivity (**a**), pH in $H_2O$ (**b**), pH in $CaCl_2$ (**c**), total carbon (C; **d**), total nitrogen (N; **e**), and C/N ratio (**f**) of the original donor soils and the corresponding repopulated soils after mixing donor soil inoculum with sterilized peat soil at a 9:1 ratio (recipient soil: donor soil). Points represent biological replicates. Original donor soils included Spanish inocula (mSP1-mSP3), and Swedish inocula (mSW1-mSW3). Sterilized peat soil is control (mC). Corresponding repopulated soils included Spanish inocula (R_mSP1-mSP3), and Swedish inocula (R_mSW1-mSW3). Boxplots show medians, interquartile ranges, and minimum-maximum values; points represent individual plants. Different letters indicate significant differences among groups based on ANOVA followed by Tukey's HSD test ($P < 0.05$).

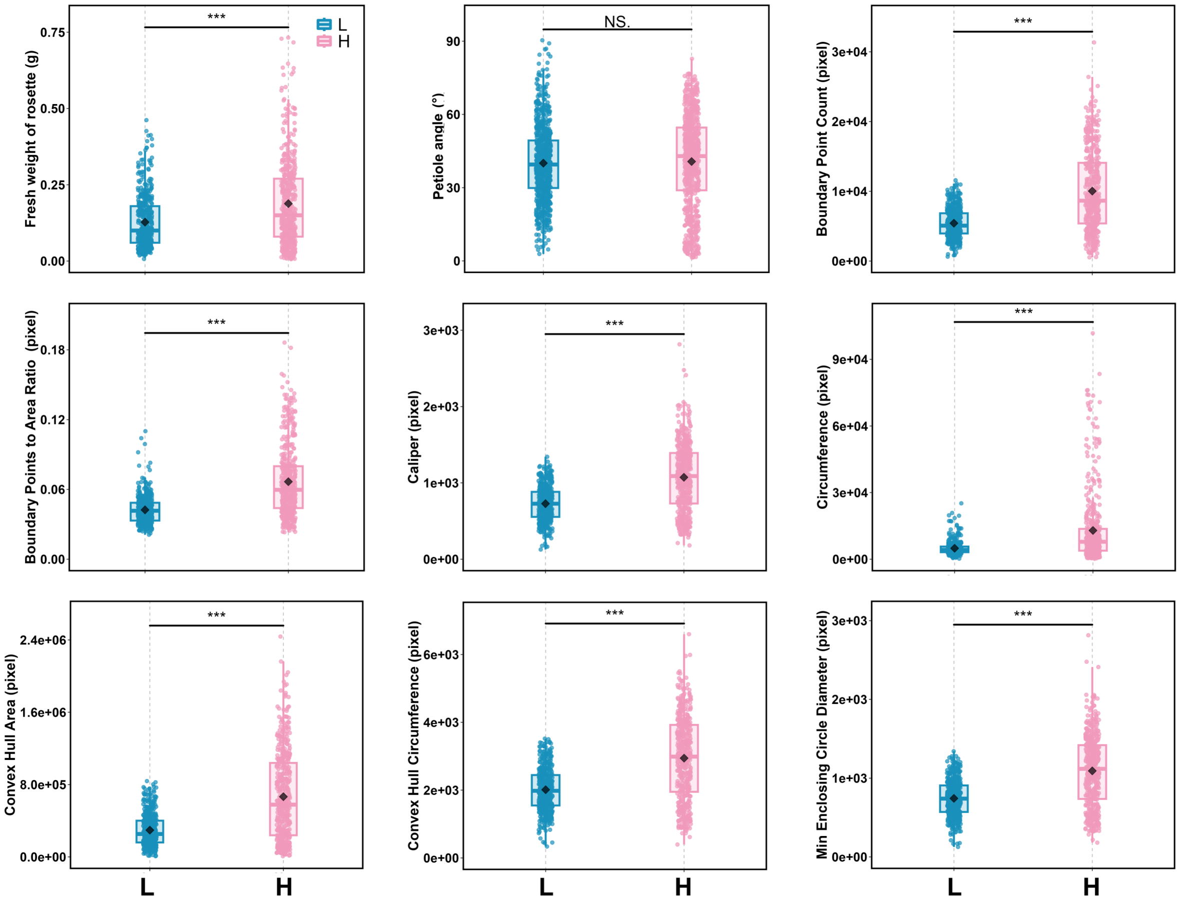


**Fig. S4. Effects of temperature on other recorded plant traits.** Temperature levels were 16 °C (L) and 28 °C (H). Other recorded plant traits measured at 14 days across the six wild accessions and *Col-0* under the two temperature treatments. Boxplots show medians, interquartile ranges, and minimum-maximum values; points represent individual plants. Asterisks denote significance between temperatures after Benjamini-Hochberg correction (*FDR < 0.05, **FDR < 0.01, ***FDR < 0.001). The plant trait analyses reported here were based on endpoint measurements taken after 14 days of temperature treatment.

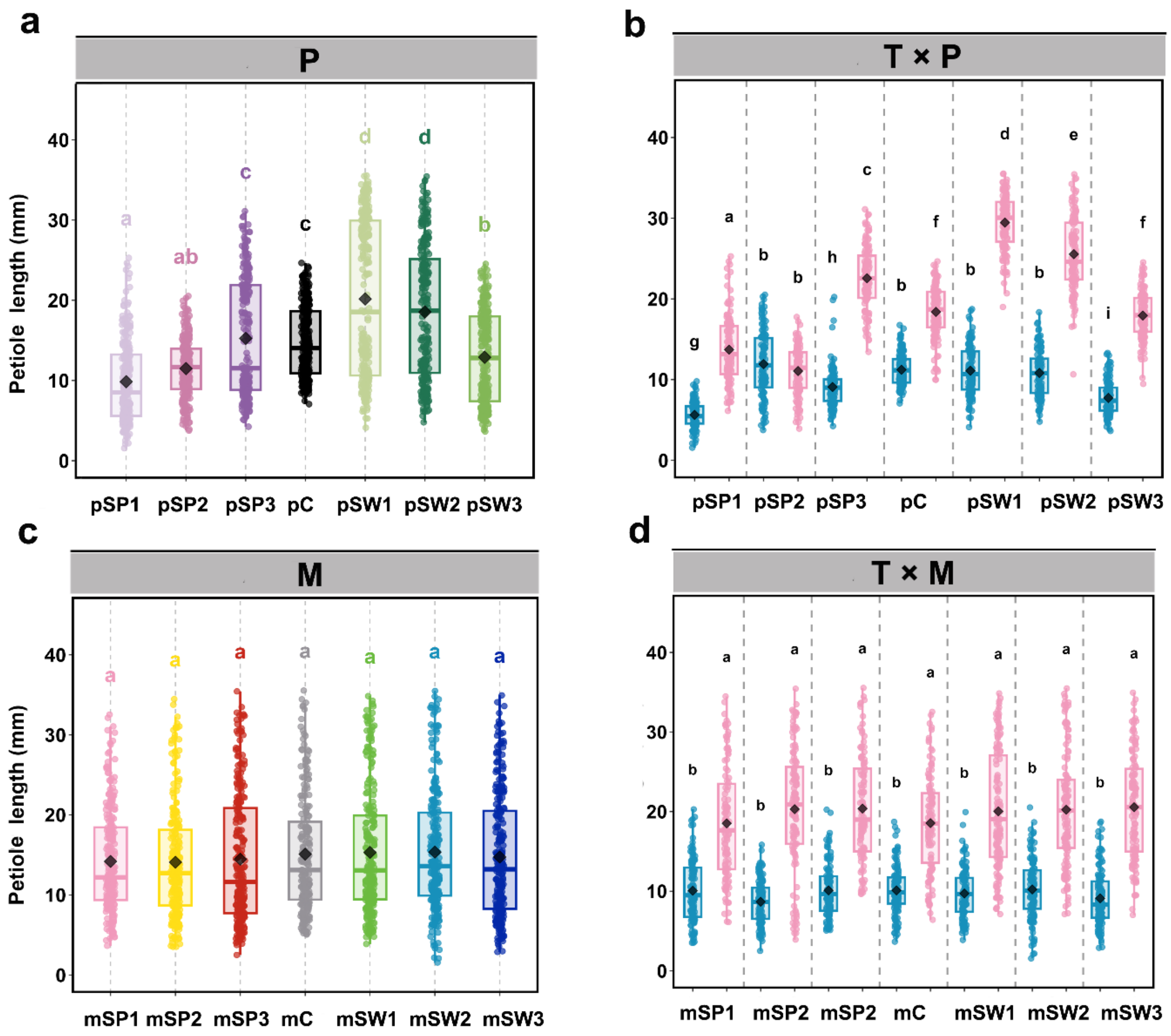


**Fig. S5. Effects of temperature, microbial inoculum, and plant genotype on petiole length.** (**a, c**) Main effects of plant genotype (P; **a**), and microbial inoculum (M; **c**) on petiole length. Temperature levels were 16 °C (L) and 28 °C (H). Microbial inoculum treatments included sterilized peat soil control (mC), Spanish inocula (mSP1-mSP3), and Swedish inocula (mSW1-mSW3). Plant genotypes included the control genotype (pC), Spanish genotypes (pSP1-pSP3), and Swedish genotypes (pSW1-pSW3). (**b, d**) Two-way interaction effects for T × P (**b**) and T × M (**d**). Boxplots show medians, interquartile ranges, and minimum-maximum values; points represent individual plants. Different letters indicate significant differences among groups based on ANOVA followed by Tukey's HSD test ($P < 0.05$). The plant trait analyses reported here were based on endpoint measurements taken after 14 days of temperature treatment.

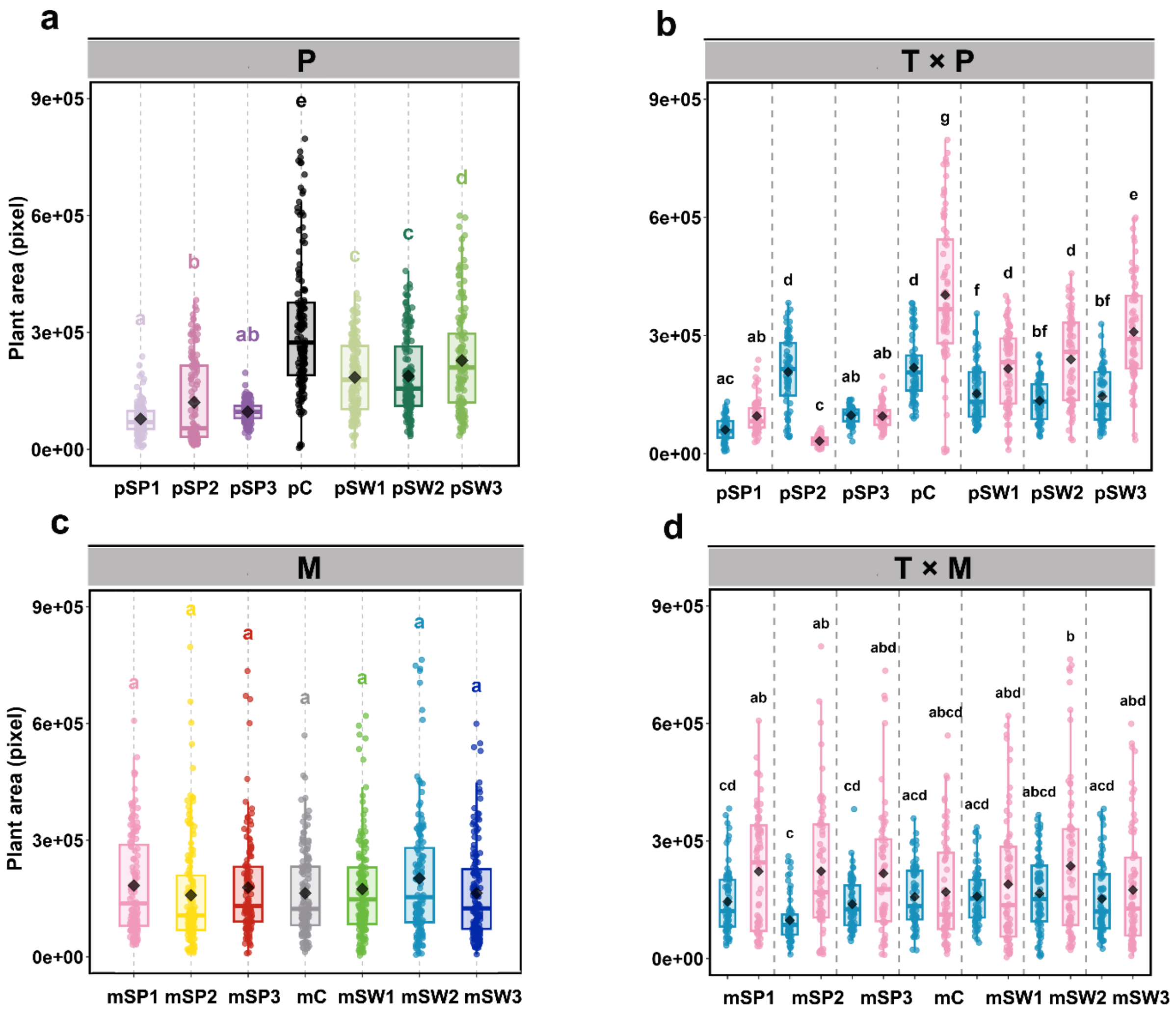


**Fig. S6. Effects of temperature, microbial inoculum, and plant genotype on plant area.** (**a, c**) Main effects of plant genotype (P; **a**), and microbial inoculum (M; **c**) on plant area. Temperature levels were 16 °C (L) and 28 °C (H). Microbial inoculum treatments included sterilized peat soil control (mC), Spanish inocula (mSP1-mSP3), and Swedish inocula (mSW1-mSW3). Plant genotypes included the control genotype (pC), Spanish genotypes (pSP1-pSP3), and Swedish genotypes (pSW1-pSW3). (**b, d**) Two-way interaction effects for T × P (**b**) and T × M (**d**). Boxplots show medians, interquartile ranges, and minimum-maximum values; points represent individual plants. Different letters indicate significant differences among groups based on ANOVA followed by Tukey's HSD test ($P < 0.05$). The plant trait analyses reported here were based on endpoint measurements taken after 14 days of temperature treatment.

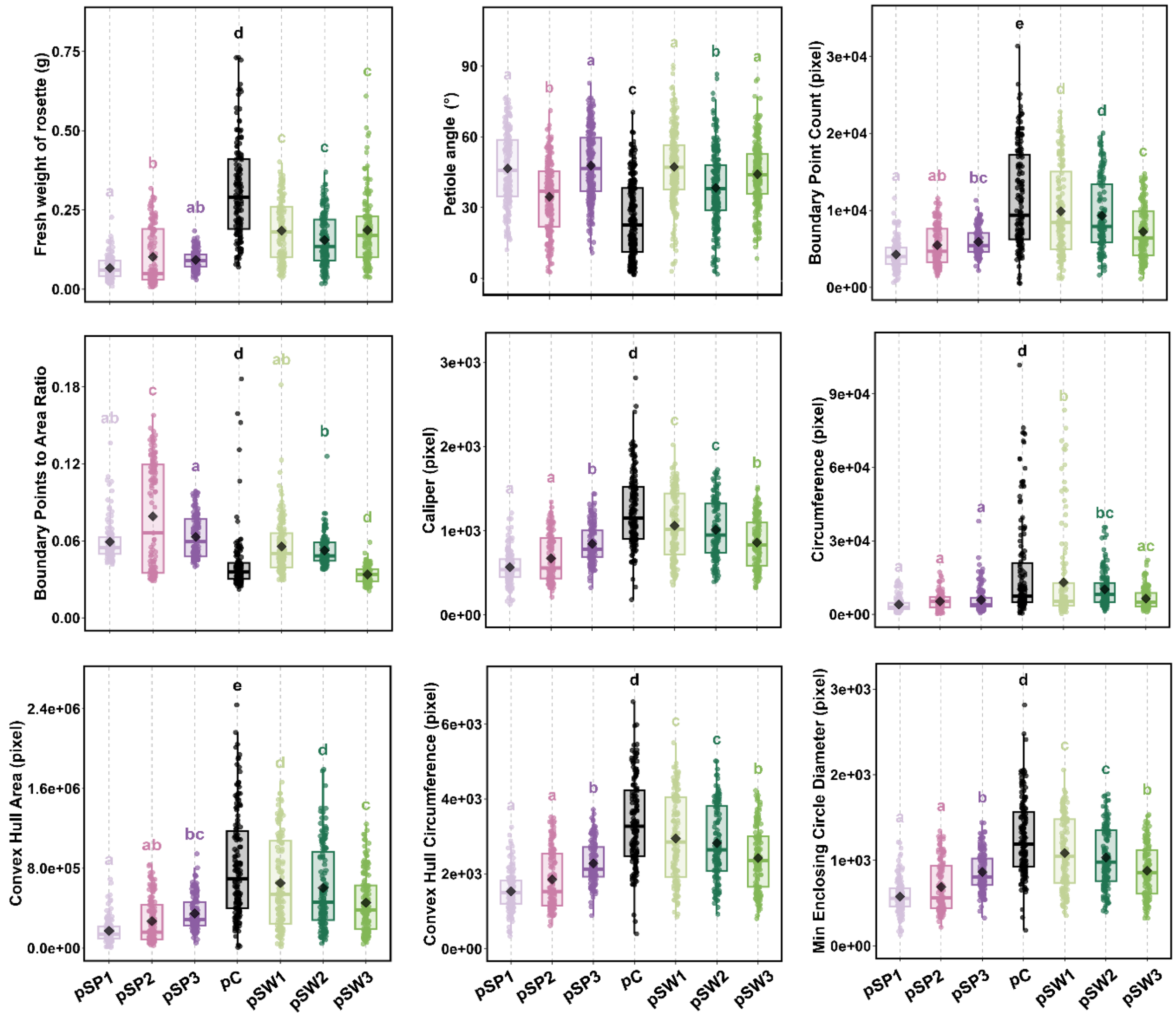


**Fig. S7. Effect of plant genotype on other recorded plant traits.** Plant genotypes included the control genotype (pC), Spanish genotypes (pSP1-pSP3), and Swedish genotypes (pSW1-pSW3). Boxplots show medians, interquartile ranges, and minimum-maximum values; points represent individual plants. Different letters indicate significant differences among groups based on ANOVA followed by Tukey's HSD test ($P < 0.05$). The plant trait analyses reported here were based on endpoint measurements taken after 14 days of temperature treatment.

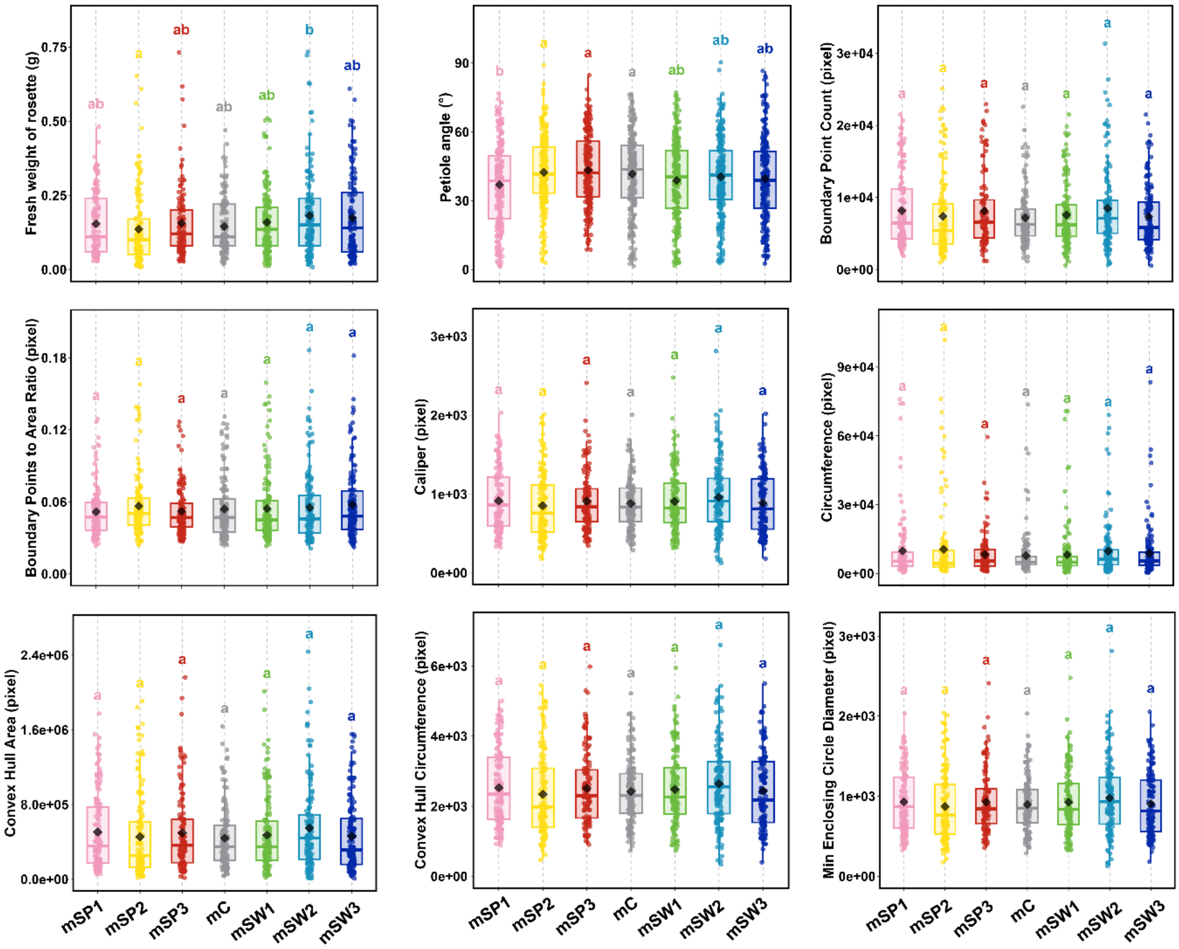


**Fig. S8 Effect of microbial inoculum on other recorded plant traits.** Microbial inoculum treatments included sterilized peat soil control (mC), Spanish inocula (mSP1-mSP3), and Swedish inocula (mSW1-mSW3). Boxplots show medians, interquartile ranges, and minimum-maximum values; points represent individual plants. Different letters indicate significant differences among groups based on ANOVA followed by Tukey's HSD test ($P < 0.05$). The plant trait analyses reported here were based on endpoint measurements taken after 14 days of temperature treatment.

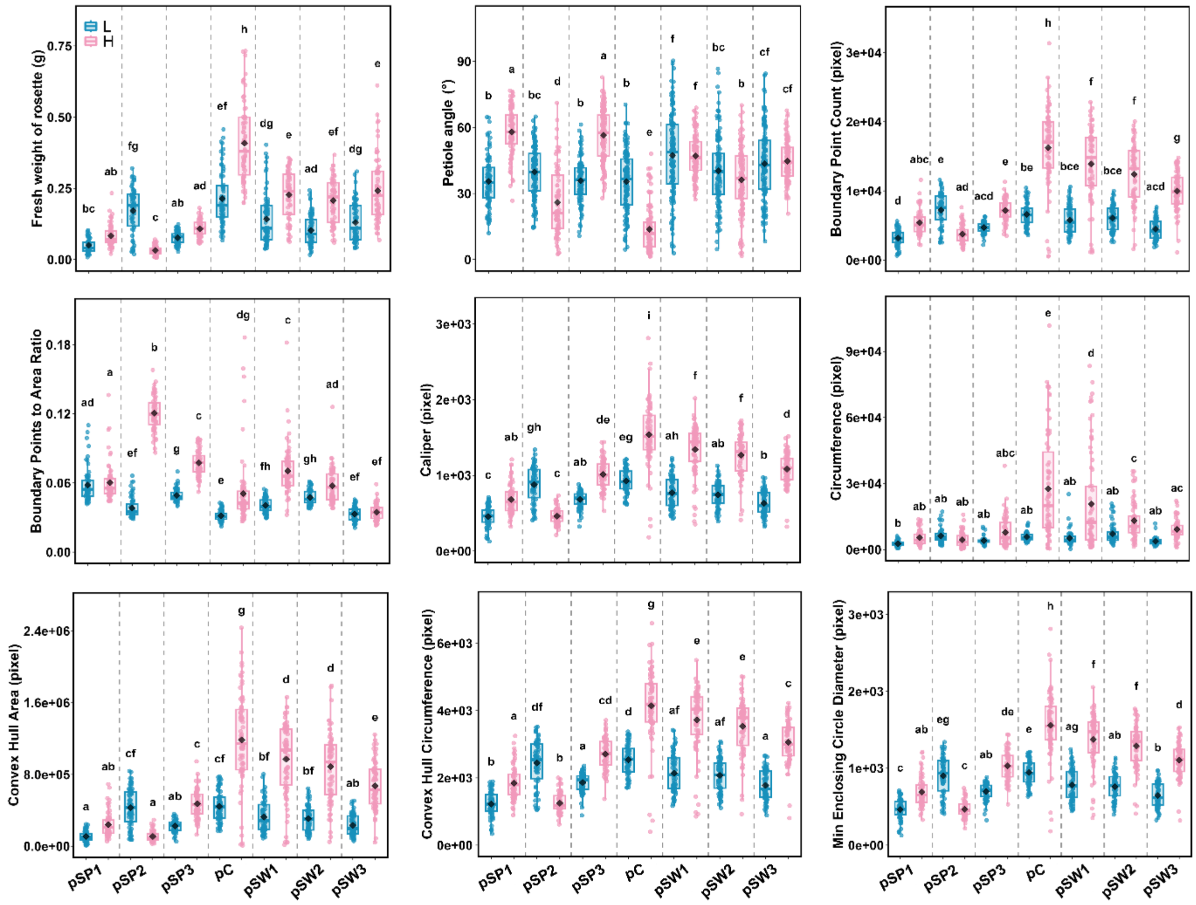


**Fig. S9. Two-way interaction effects for temperature and plant genotype (T × P) on other recorded plant traits.** Temperature levels were 16 °C (L) and 28 °C (H). Plant genotypes included the control genotype (pC), Spanish genotypes (pSP1-pSP3), and Swedish genotypes (pSW1-pSW3). Boxplots show medians, interquartile ranges, and minimum-maximum values; points represent individual plants. Different letters indicate significant differences among groups based on ANOVA followed by Tukey's HSD test ($P < 0.05$). The plant trait analyses reported here were based on endpoint measurements taken after 14 days of temperature treatment.

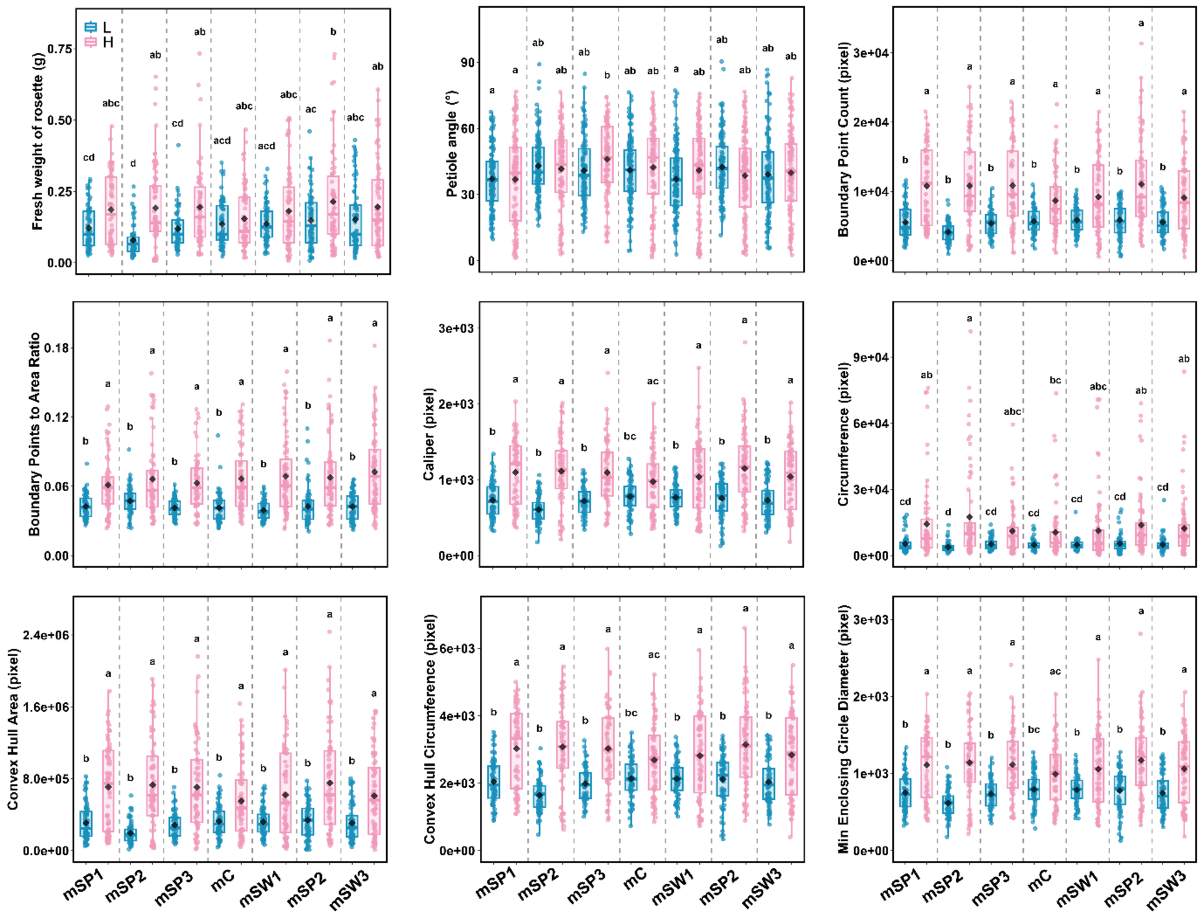

**Fig. S10. Two-way interaction effects for temperature and microbial inoculum (T × M) on other recorded plant traits.** Temperature levels were 16 °C (L) and 28 °C (H). Microbial inoculum treatments included sterilized peat soil control (mC), Spanish inocula (mSP1-mSP3), and Swedish inocula (mSW1-mSW3). Boxplots show medians, interquartile ranges, and minimum-maximum values; points represent individual plants. Different letters indicate significant differences among groups based on ANOVA followed by Tukey's HSD test ($P$ < 0.05). The plant trait analyses reported here were based on endpoint measurements taken after 14 days of temperature treatment.

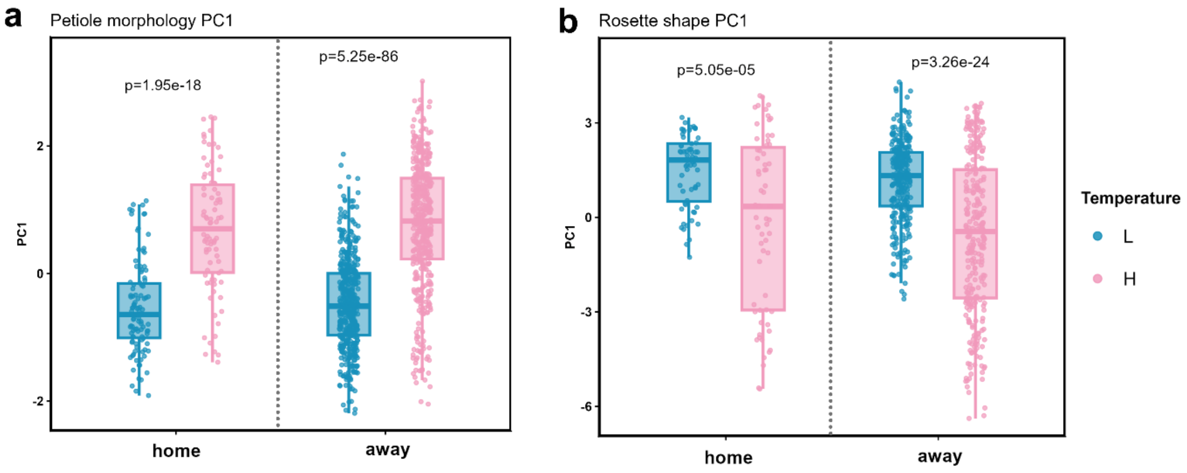


**Fig. S11. Principal component summaries of temperature-dependent variation in plant morphology.** (**a**) Distribution of PC1 scores from the PCA of petiole-related traits, including petiole angle and petiole length. (**b**) Distribution of PC1 scores from the PCA of rosette shape traits measured with the LemnaTec phenotyping platform. PC1 scores summarize coordinated multivariate responses of plant architecture to temperature. Boxplots show medians, interquartile ranges, and minimum-maximum values; points represent individual plants. *P* values between temperature treatments after Benjamini-Hochberg correction (FDR < 0.05) were shown in the plots.

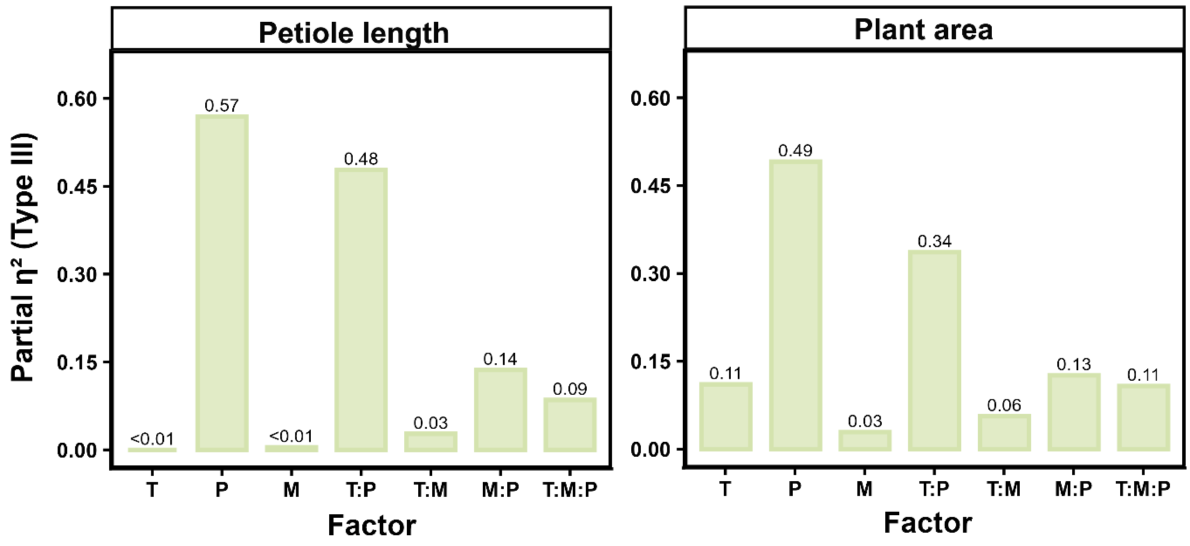


**Fig. S12. Variance partitioning of plant phenotypic traits under the factorial T × M × P design.** Partial $\eta^2$ values derived from Type III ANOVA are shown for plant area and petiole length. Models included temperature (T), microbial inoculum (M), plant genotype (P), and their interaction terms. Effect sizes represent the proportion of trait variance attributable to each factor after accounting for the other model terms. For petiole length, the temperature main effect was absorbed by strong genotype-dependent interactions (T × P), resulting in no estimable uniform temperature effect under the Type III framework. Values above bars indicate partial $\eta^2$.

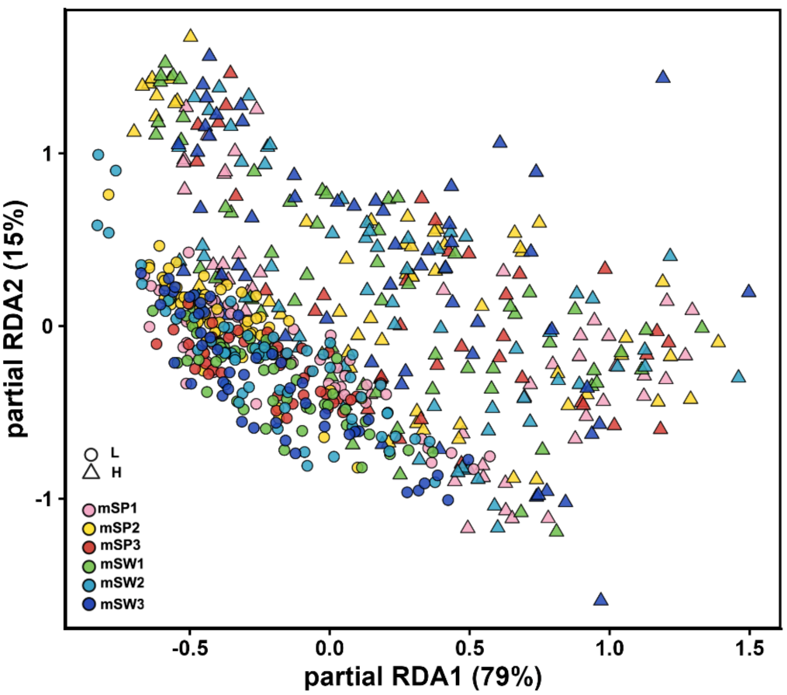


**Fig. S13. Partial redundancy analysis (RDA) of plant trait variation after conditioning on soil chemistry.** Partial RDA was performed on scaled multivariate plant trait data using temperature (T), inoculum identity (M), and plant genotype (P) as explanatory variables, while conditioning on soil nitrogen and soil C/N ratio. Points represent individual plant samples. Fill colors indicate inoculum identity, and shapes indicate temperature treatment, with circles representing low temperature and triangles representing high temperature. Axes show the proportion of constrained variation represented by each partial RDA axis after accounting for soil N and C/N. Term significance was assessed using permutation tests with 999 permutations.

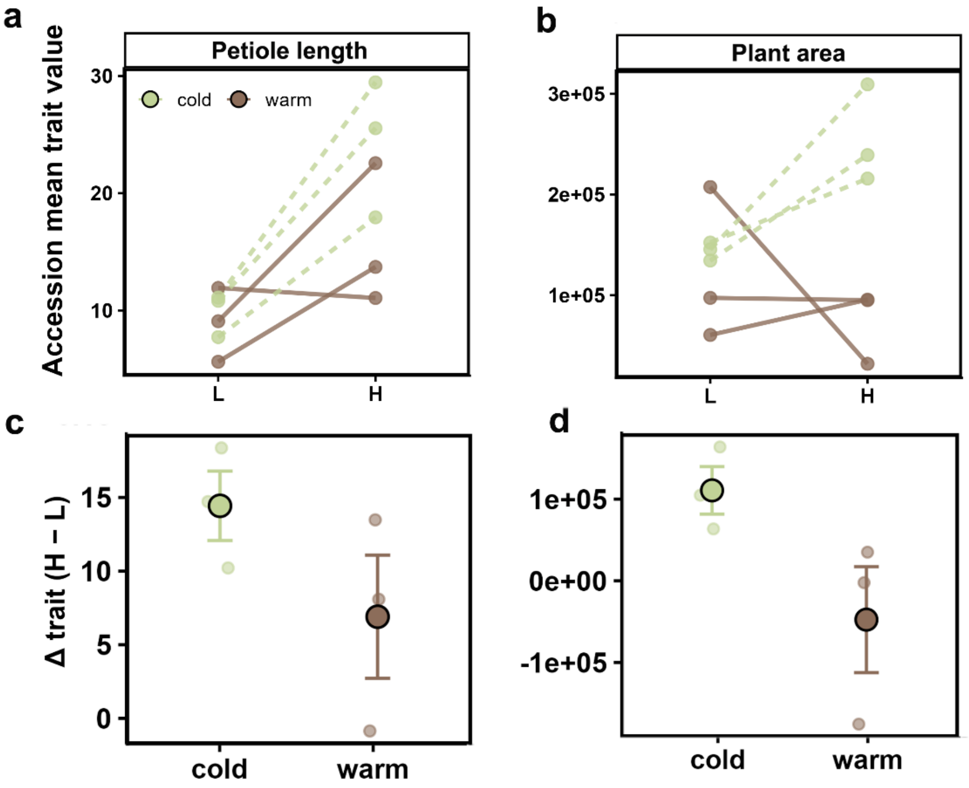


**Fig. S14. Temperature-induced plasticity differs between climatic origins.** (**a, b**) Accession-level mean values of petiole length (**a**) and plant area (**b**) across the six wild accessions under low (L, 16 °C) and high (H, 28 °C) temperature treatments. Each line represents one accession; solid lines indicate warm-origin accessions and dashed lines indicate cold-origin accessions. Points represent accession means across biological replicates. (**c, d**) Distributions of accession-level plastic responses quantified as Δ(H–L) for petiole length (**c**) and plant area (**d**). Positive values indicate increased trait expression at elevated temperature. Small points represent individual accessions. Larger symbols indicate group means, with error bars showing ± SE.

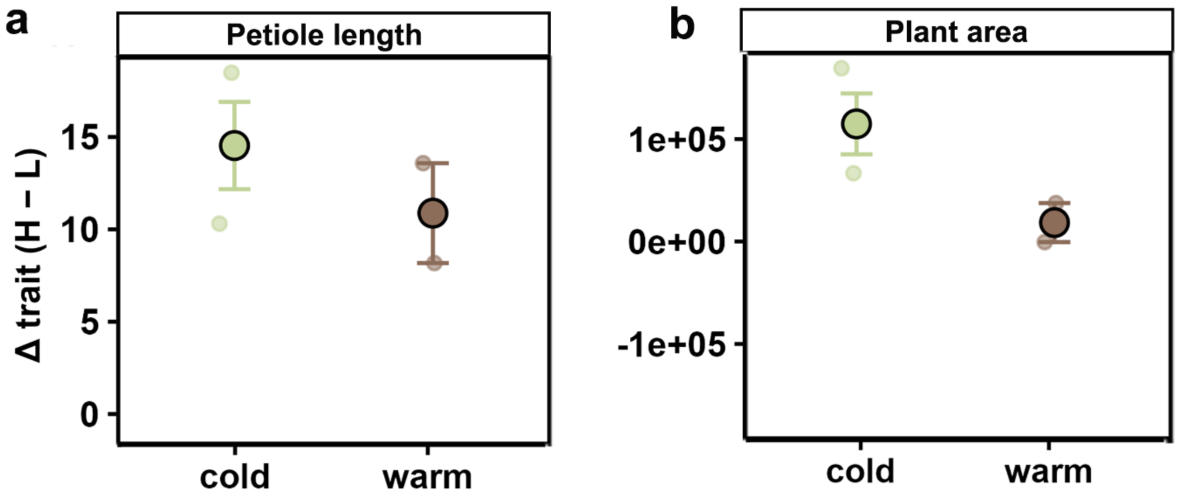


**Fig. S15. Sensitivity analysis excluding accession IP-EVS-0_B.** Comparison of temperature-response values, calculated as Δ(H–L), for the analyzed plant traits a reduced dataset excluding the warm-origin accession IP-EVS-0_B. Small points represent individual accessions. Larger symbols indicate group means, with error bars showing ± SE. This analysis was used to assess whether the climatic-origin comparison was disproportionately influenced by IP-EVS-0_B, which showed marked growth suppression under elevated temperature.

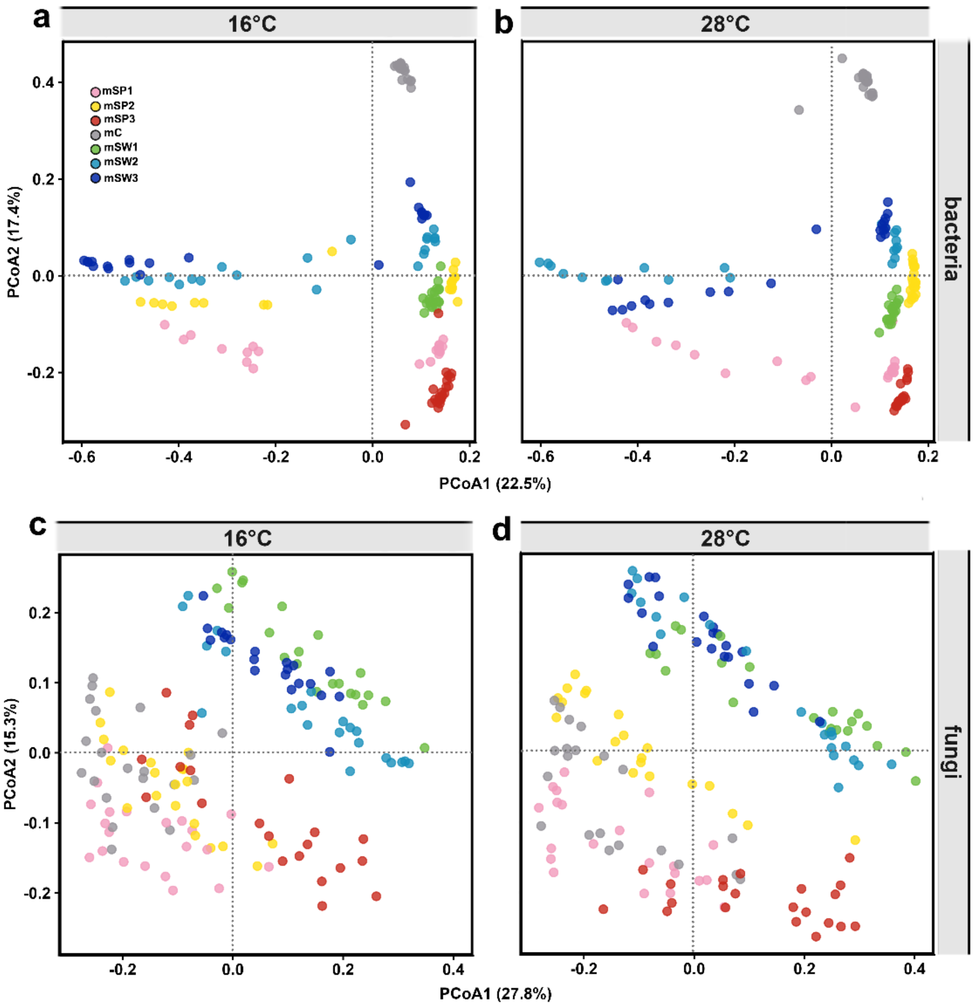


**Fig. S16. Temperature-stratified ordination of soil-associated rhizosphere community structure.** Principal coordinate analysis (PCoA) of bacterial (**a, b**) and fungal (**c, d**) rhizosphere communities based on Bray-Curtis dissimilarity, stratified by temperature treatment (16 °C and 28 °C). Points represent individual samples and are colored by microbial inoculum origin.

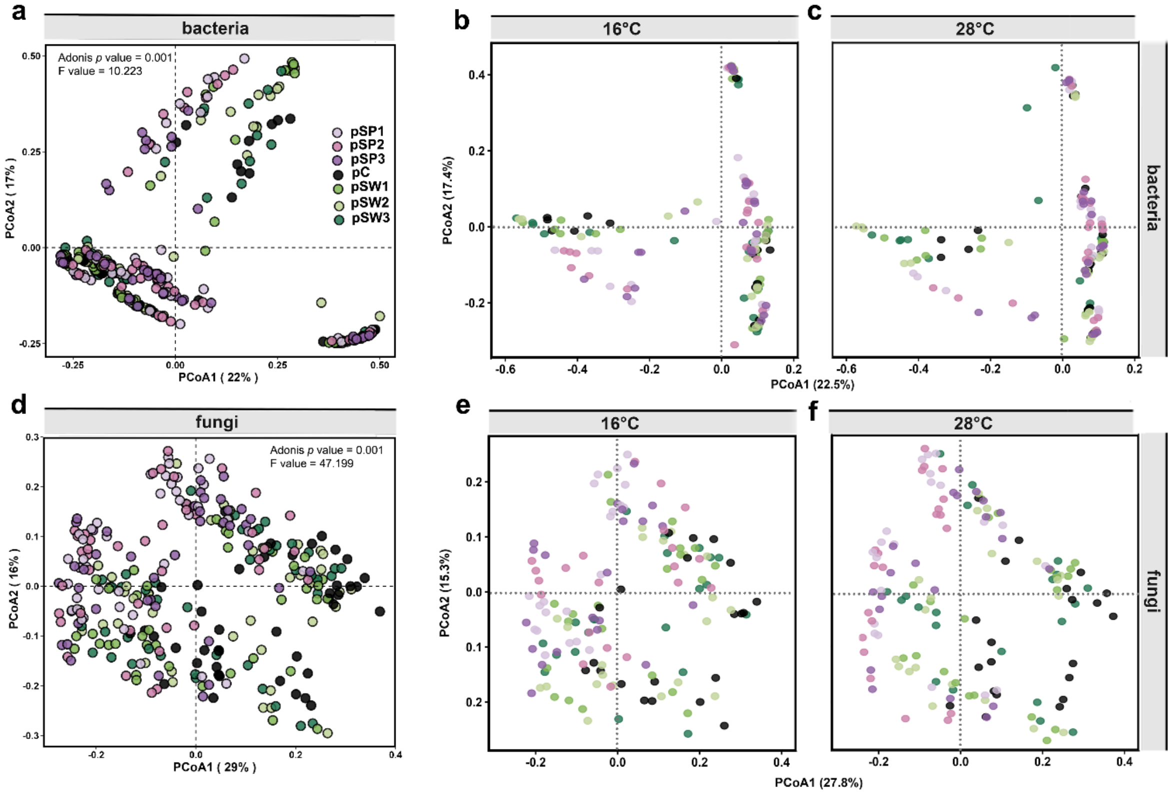


**Fig. S17. Temperature-stratified ordination of plant-associated rhizosphere community structure.** Principal coordinate analysis (PCoA) of bacterial (**a-c**) and fungal (**d-f**) rhizosphere communities based on Bray-Curtis dissimilarity, stratified by temperature treatment (16 °C and 28 °C). Points represent individual samples and are colored by plant genotype.

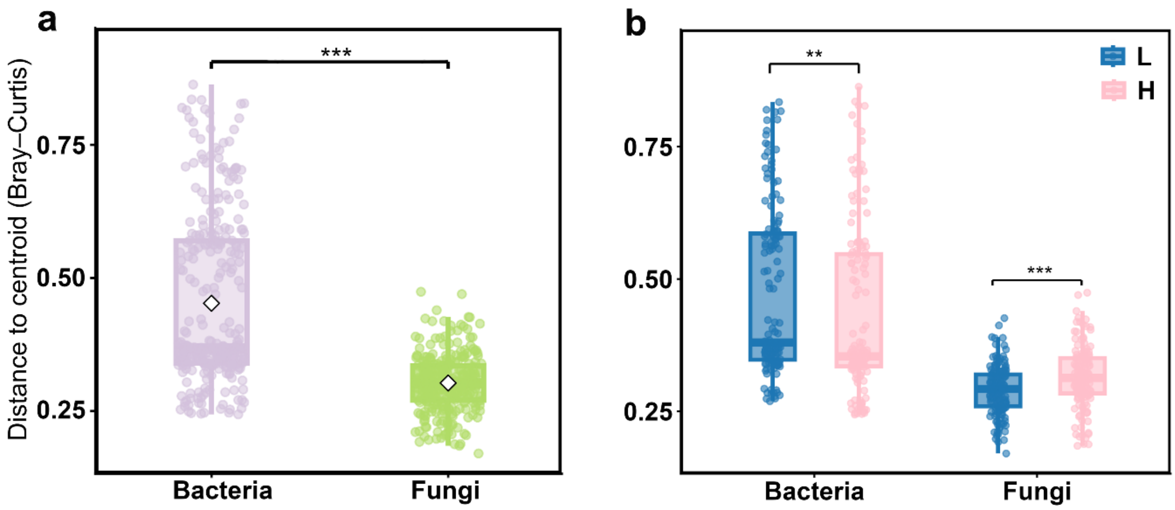


**Fig. S18. (a) Multivariate dispersion of rhizosphere bacterial and fungal communities. Dispersion was compared between microbial kingdoms. (b) Temperature-dependent multivariate dispersion of rhizosphere bacterial and fungal communities.** Dispersion was compared between low (L, 16 °C) and high (H, 28 °C) temperature treatments within each microbial kingdom. Multivariate dispersion was calculated as the distance of each sample to its group centroid in principal coordinate space based on Bray-Curtis dissimilarity using betadisper in the vegan package (999 permutations). Boxplots show medians and interquartile ranges; points represent individual samples. Asterisks denote significance between temperatures after Benjamini-Hochberg correction (*FDR < 0.05, **FDR < 0.01, ***FDR < 0.001).

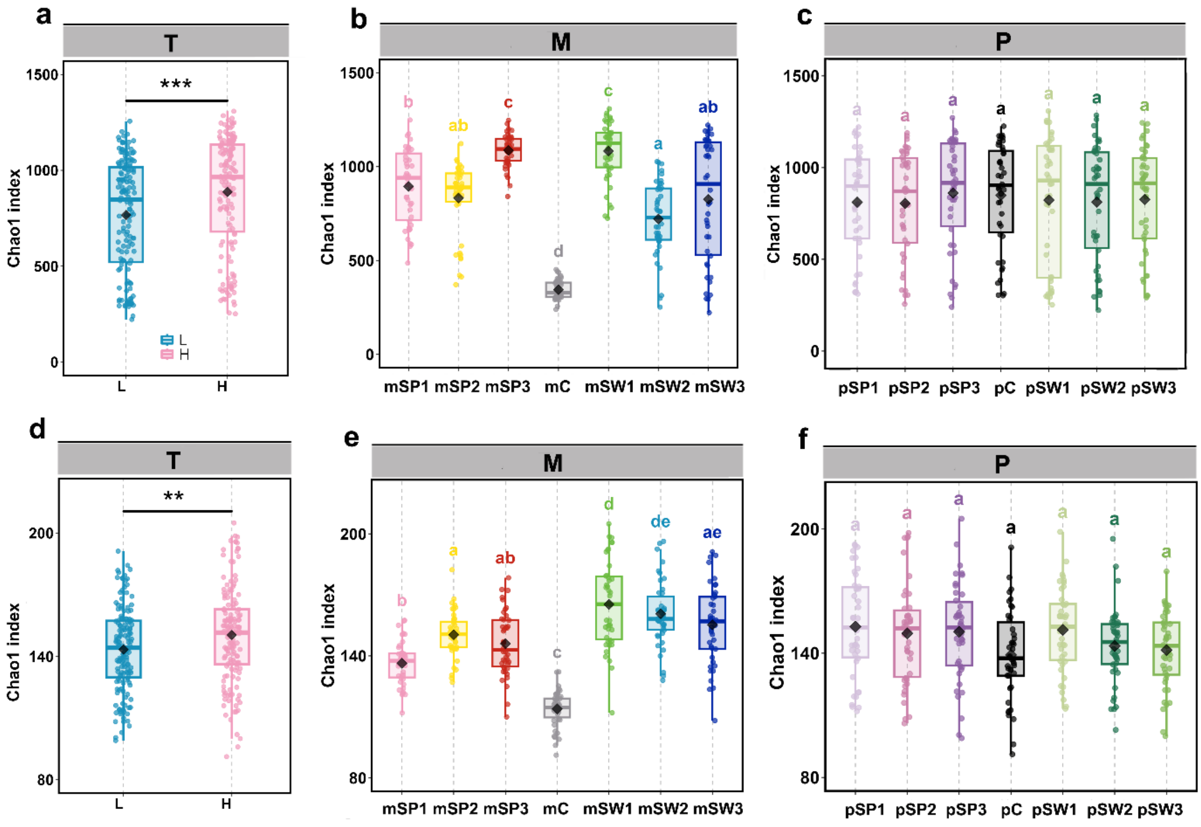


**Fig. S19. Alpha diversity of rhizosphere bacterial and fungal communities across temperature, microbial inoculum, and plant genotype treatments.** Alpha-diversity metrics for bacterial (**a-c**) and fungal (**d-f**) rhizosphere communities across the factorial combinations of temperature (T), microbial inoculum (M), and plant genotype (P). Temperature levels were 16 °C (L) and 28 °C (H). Microbial inoculum treatments included sterilized peat soil control (mC), Spanish inocula (mSP1-mSP3), and Swedish inocula (mSW1-mSW3). Plant genotypes included the control genotype (pC), Spanish genotypes (pSP1-pSP3), and Swedish genotypes (pSW1-pSW3). Panels show the diversity metric(s) used in the main analysis for each microbial kingdom. Boxplots show medians, interquartile ranges, and minimum-maximum values; points represent individual samples. Asterisks denote significance between temperatures after Benjamini-Hochberg correction (*FDR < 0.05, **FDR < 0.01, ***FDR < 0.001). Different letters indicate significant differences among groups based on ANOVA followed by Tukey's HSD test ($P < 0.05$).

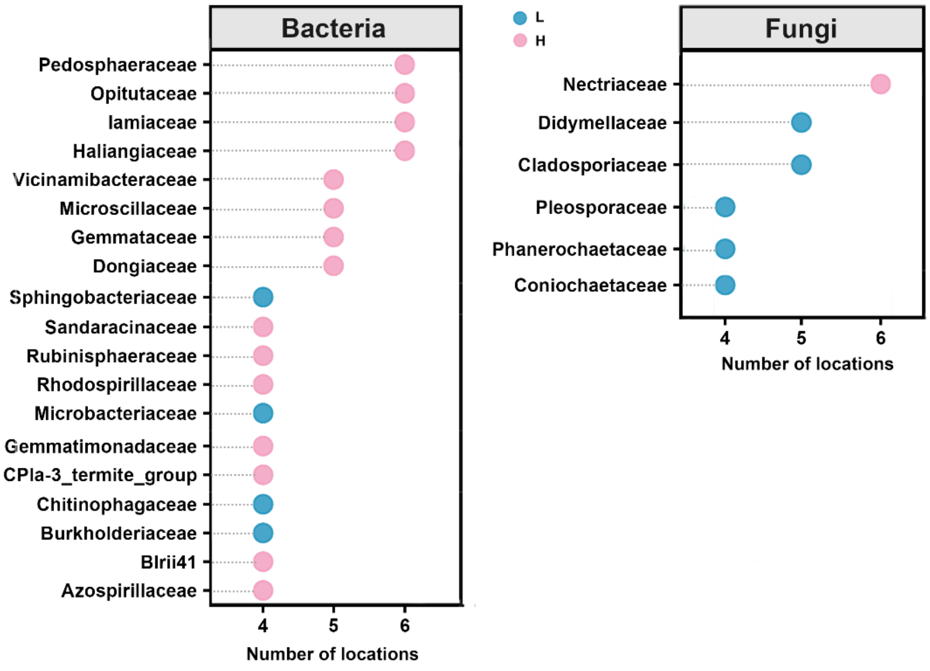


**Fig. S20. Conserved temperature-associated microbial taxa across locations.** Bacterial (left) and fungal (right) families that were repeatedly identified as significantly associated with temperature in four or more sampling sites. The x-axis indicates the number of locations in which each family was detected, and colors indicate the direction of association, with enrichment under 16 °C (L) and 28 °C (H).

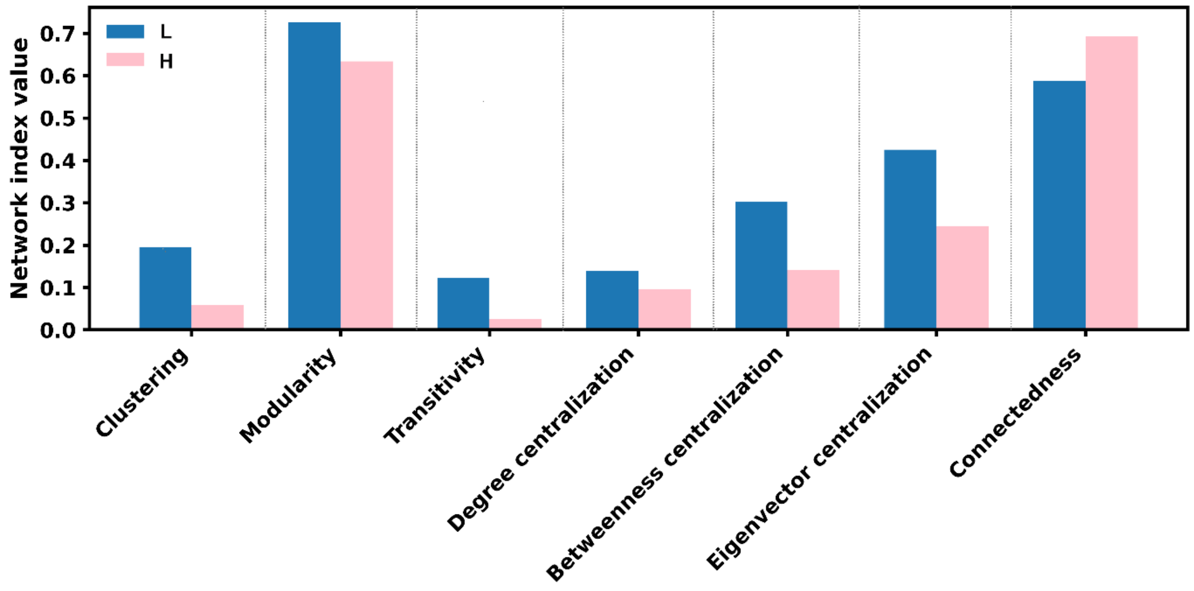


**Fig. S21. Comparison of network topological properties between the 16 °C and 28 °C rhizosphere microbial co-occurrence networks.** Network topological indices calculated for the temperature-specific microbial co-occurrence networks shown in **Fig. 5**. Parameters include measures of connectedness and centralization, such as clustering coefficient, transitivity, modularity, betweenness centrality, and eigenvector centralization, as computed in the Molecular Ecological Network Analysis Pipeline (MENAP). Bars show the values for the 16 °C and 28 °C network.

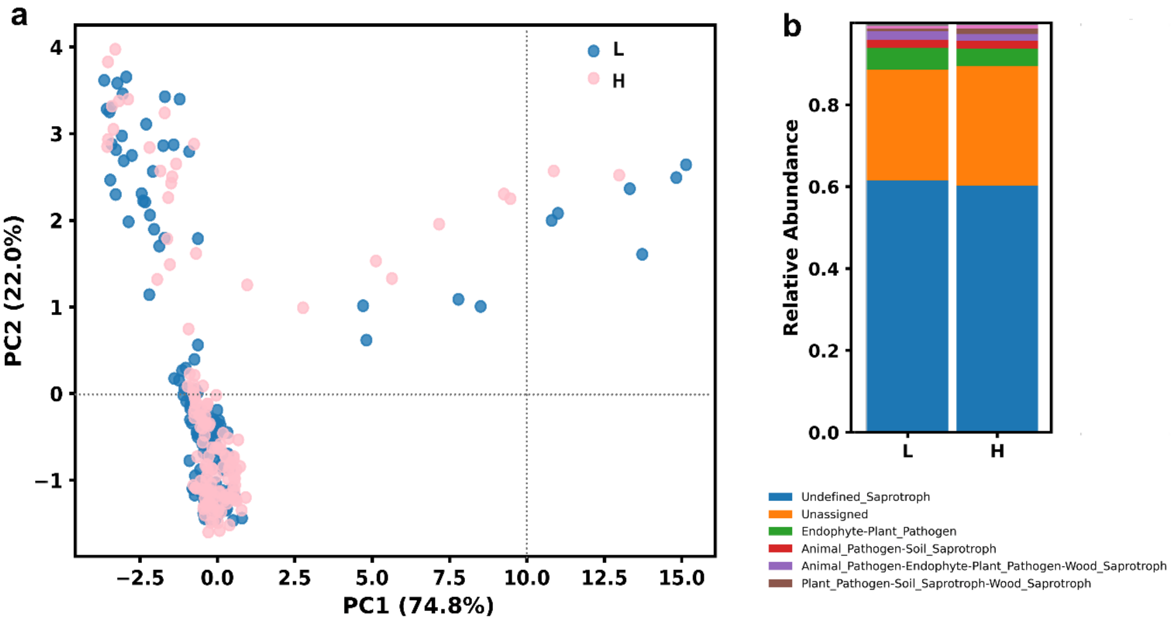


**Fig. S22. Predicted functional profiles of rhizosphere microbial communities under moderate warming.** (**a**) Principal component analysis (PCA) of predicted bacterial metabolic pathways inferred using PICRUSt2 for rhizosphere bacterial communities at 16 °C (L) and 28 °C (H). (**b**) Fungal trophic guild composition inferred using FUNGuild across the two temperature treatments.

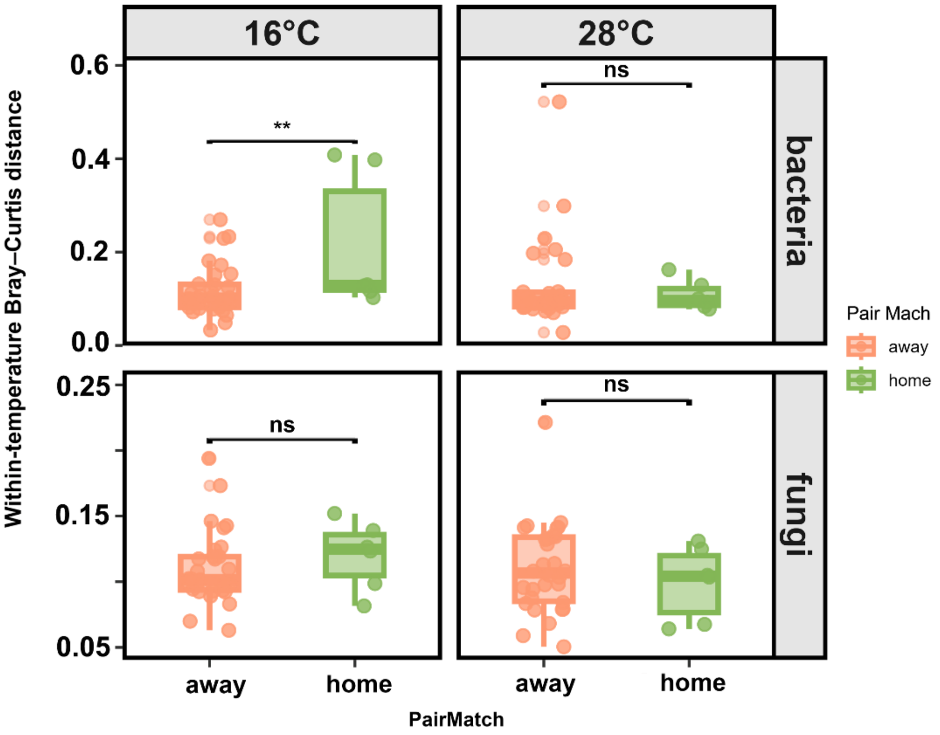


**Fig. S23. Rhizosphere microbiome turnover under home and away conditions across temperatures.** Rhizosphere microbiome turnover (Δβ), quantified as Bray-Curtis dissimilarity, under home and away microbiome conditions at 16 °C and 28 °C for bacterial and fungal communities. Panels show bacterial communities at 16 °C (upper left) and 28 °C (upper right), and fungal communities at 16 °C (lower left) and 28 °C (lower right). Home conditions refer to plant-microbial inoculum combinations originating from the same site; away conditions refer to combinations from different sites. Differences between home and away conditions within each panel were assessed using Wilcoxon rank-sum tests. Points represent individual samples, and boxes indicate medians and interquartile ranges. Asterisks denote significance after Benjamini-Hochberg correction (FDR < 0.05, FDR < 0.01, FDR < 0.001).

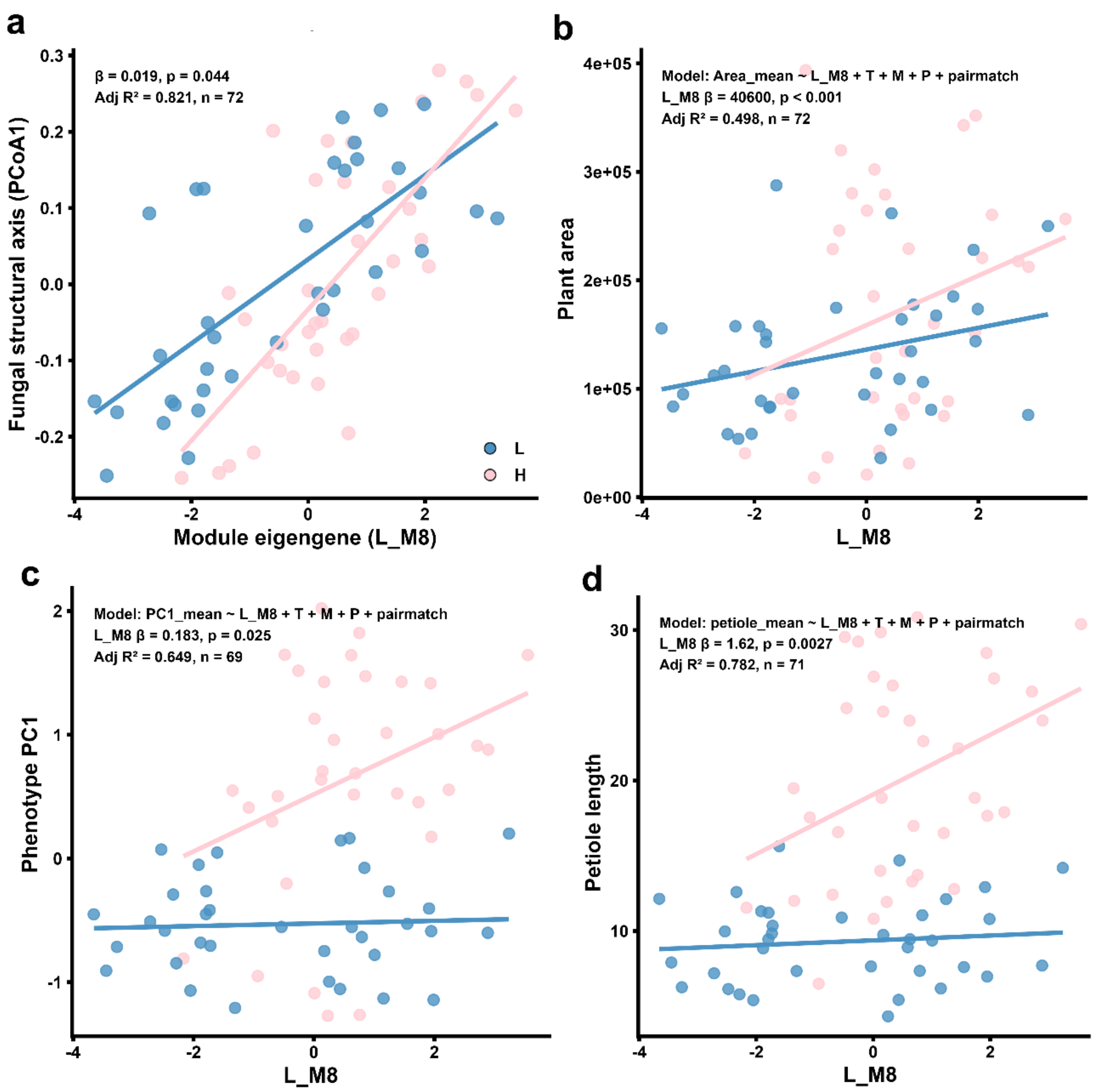


**Fig. S24. Network module eigengene L_M8 links local interaction structure to plant phenotype and fungal community organization.** Relationships involving the 16 °C network module eigengene L_M8, the only module retained as a significant module-level predictor in the regression analysis. Panels show associations between L_M8 and fungal community PCoA1(**a**), plant area (**b**), phenotype PC1 (**c**), and petiole length (**d**). Each point represents one aggregated treatment combination. Lines indicate fitted linear regressions. Temperature levels were 16 °C (L) and 28 °C (H). Statistical annotations report the corresponding model fit and significance for each relationship. No module-level predictors were retained from the 28 °C network.

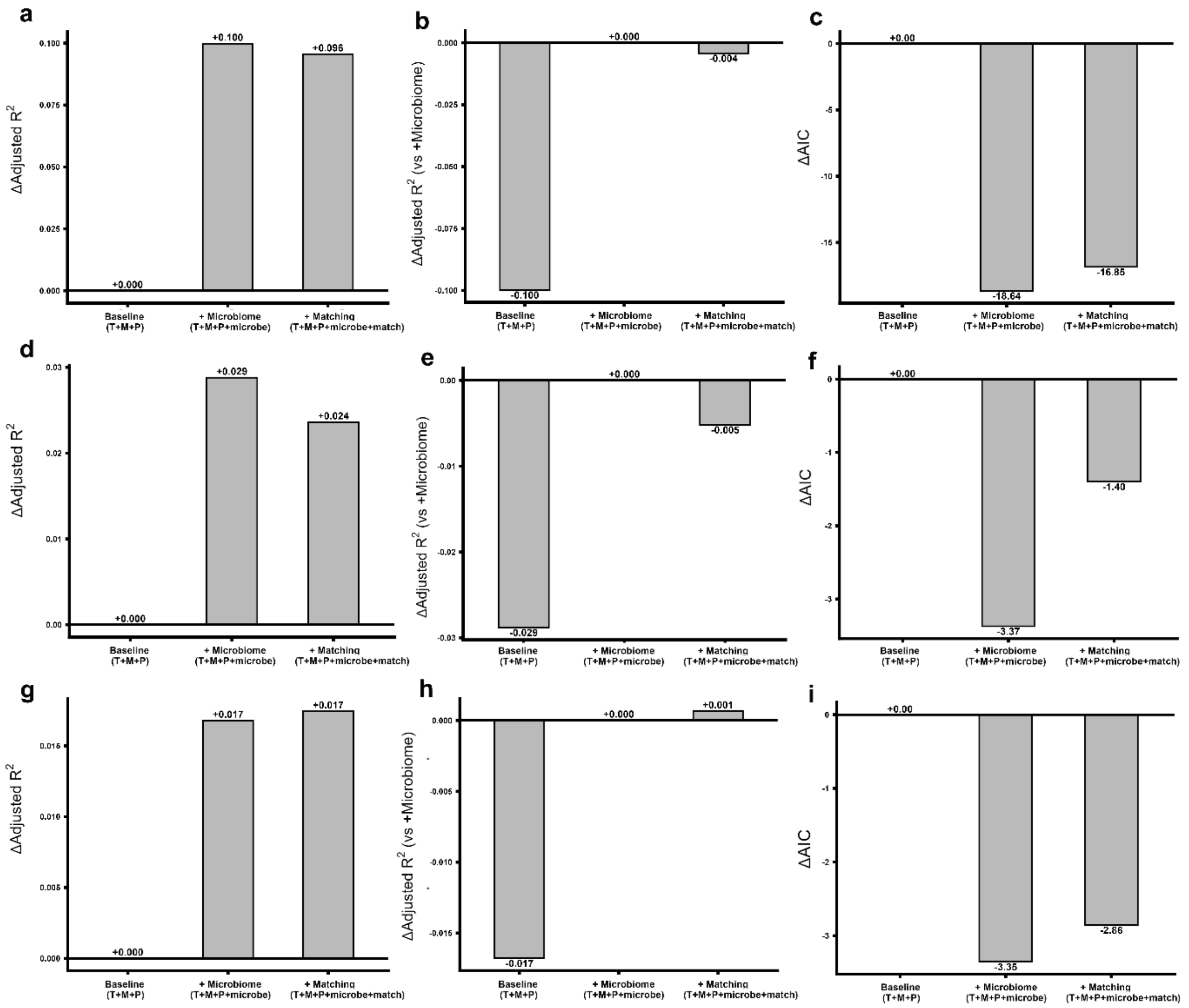


**Fig. S25. Incremental model performance gains across plant traits.** Changes in adjusted $R^2$ and AIC across hierarchical linear models predicting plant area (**a-c**), phenotype PC1 (**d-f**), and petiole length (**g-i**). The baseline model included temperature (T), microbial inoculum (M), and plant genotype (P). The second model additionally included microbial composition-derived predictors (fungal and bacterial Δβ and PCoA1), and the third model further included host-microbiome matching. Panels show Δadjusted $R^2$ relative to the baseline model (**a, d, g**), Δadjusted $R^2$ relative to model 2 (**b, e, h**), and ΔAIC relative to the baseline model (**c, f, i**).

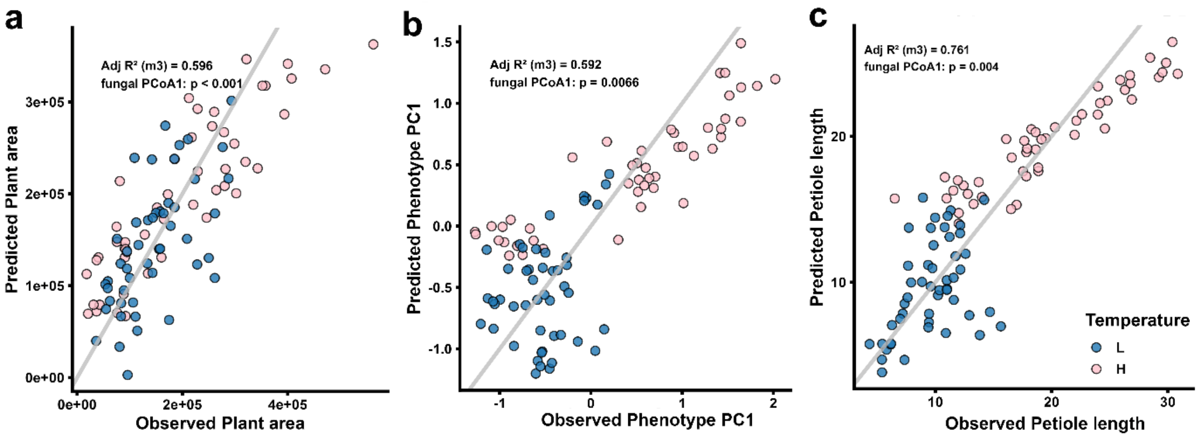


**Fig. S26. Observed-versus-predicted relationships for fungal-structure-based models of plant traits.** Observed versus predicted values for plant area (**a**), phenotype PC1 (**b**), and petiole length (**c**) derived from linear models including temperature (T), microbial inoculum (M), plant genotype (P), host-microbiome matching, and fungal microbiome composition-derived predictors (fungal PCoA1 and Δβ). The diagonal line represents the 1:1 expectation. Each point represents a treatment-level aggregate corresponding to one T × M × P combination.

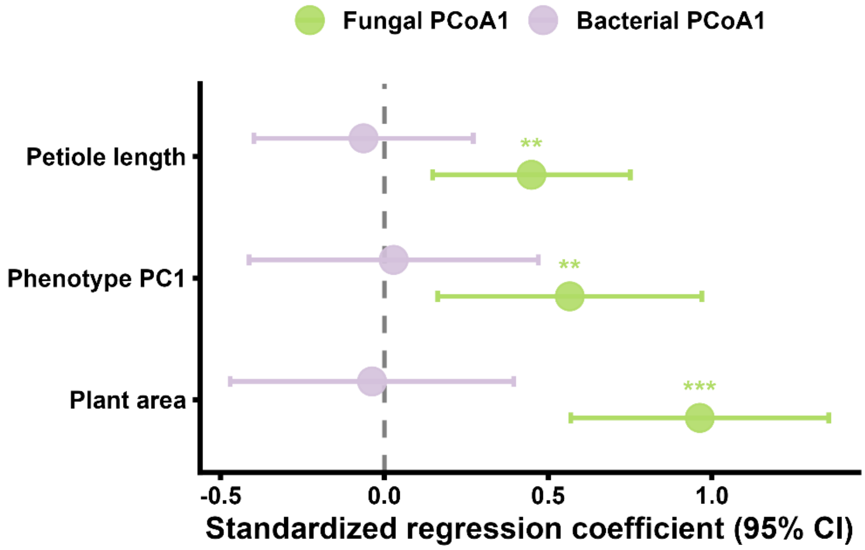


**Fig. S27. Standardized regression coefficients of fungal and bacterial community composition across plant traits.** Forest plots showing standardized regression coefficients (±95% confidence intervals) for fungal PCoA1 and bacterial PCoA1 from linear models fitted separately for plant area, phenotype PC1, and petiole length. All continuous predictors and response variables were standardized before model fitting to allow direct comparison of effect sizes across traits. Models included temperature (T), microbial inoculum (M), plant genotype (P), host-microbiome matching, and microbiome turnover metrics (fungal and bacterial Δβ) as covariates. Error bars denote 95% confidence intervals.
